# DypFISH: Dynamic Patterned FISH to Interrogate RNA and Protein Spatial and Temporal Subcellular Distribution

**DOI:** 10.1101/536383

**Authors:** Anca F. Savulescu, Robyn Brackin, Emmanuel Bouilhol, Benjamin Dartigues, Jonathan H. Warrell, Mafalda R. Pimentel, Stephane Dallongeville, Jan Schmoranzer, Jean-Christophe Olivo-Marin, Edgar R. Gomes, Macha Nikolski, Musa M. Mhlanga

**Author notes:** These authors contributed equally to this work (Anca F. Savulescu, Robyn Brackin). The authors declare that this material has not been published elsewhere in any language is not under simultaneous consideration for publication elsewhere and will not be submitted for publication elsewhere whilst under consideration.

## Abstract

Advances in single cell RNA sequencing have allowed for the identification and characterization of cellular subtypes based on quantification of the number of transcripts in each cell. However, cells may differ not only in the number of mRNA transcripts that they exhibit, but also in their spatial and temporal distribution, intrinsic to the definition of their cellular state. Here we describe DypFISH, an approach to quantitatively investigate the spatial and temporal subcellular localization of RNA and protein, by combining micropatterning of cells with fluorescence microscopy at high resolution. We introduce a range of analytical techniques for quantitatively interrogating single molecule RNA FISH data in combination with protein immunolabeling over time. Strikingly, our results show that constraining cellular architecture reduces variation in subcellular mRNA and protein distributions, allowing the characterization of their localization and dynamics with high reproducibility. Many tissues contain cells that exist in similar constrained architectures. Thus DypFISH reveals reproducible patterns of clustering, strong correlative influences of mRNA-protein localization on MTOC orientation when they are present and interdependent dynamics globally and at specific subcellular locations which can be extended to physiological systems.

## INTRODUCTION

The need to incorporate subcellular spatial information of central dogma molecules into traditional omics approaches has led to the call for spatially resolved omics of various kinds (Crosetto et al., 2015). This has become more urgent as projects such as the Human Cell Atlas begin to use technologies such as single cell RNA sequencing (scRNA seq) to characterize subtypes of cells based on their molecular signatures by counting the number of RNA transcripts. Although much progress has been made in spatially resolved transcriptomics (reviewed in Crosetto et al., 2015, Medioni and Besse, 2018, Strell et al., 2018) incorporating spatial information into omics approaches carries with it several difficulties, such as coping with biological heterogeneity and noise in the spatial domain and developing analytical approaches, which avoid the loss of spatial information. Many current models for the measurement of gene expression neglect spatial information (Raj et al., 2006, Tanaguchi et al., 2010) and are not directly applicable in contexts where expression is highly localized. Moreover, such localization may be highly indicative of cell state and not fully reflected in scRNA seq data. Indeed though scRNA seq has identified several new cell types, spatial position of RNA, which is highly influential to cell state remains unincorporated in such measurements. By revealing 3D positions of RNA, for example, cells states as defined by scRNA seq may be altered or redefined gaining higher resolution and information content.

The importance of subcellular localization of mRNA transcripts as a means to spatially and temporally restrict translation has been demonstrated in a wide variety of cell types (Bashirullah et al., 1998, Besse and Ephrussi, 2008, Jansen, 2001, Kloc et al., 2002, and Martin and Ephrussi, 2009, Zappulo et al., 2017). Localizing specific mRNA transcripts to distinct subcellular localizations therefore serves as an important determinant of protein localization and is often highly influential to cell state (Moor et al., 2017, Zappulo et al., 2017). Although the number of fully characterised localized mRNAs is currently small, emerging studies have demonstrated that the mRNA localization phenomenon is more widespread than previously assumed and may in fact be relevant for the majority of mRNA transcripts (Bouvrette et al., 2017, La Manno et al., 2018, Lecuyer et al., 2007, Moor et al., 2017, Sharp et al., 2011, Weis et al., 2013,, Zappulo et al., 2017). Indeed such evidence has extended to distinct subcellular localization patterns for cytoplasmic and nuclear localization of long noncoding RNAs (lncRNAs) has recently emerged (Cabili et al., 2015).

Studies showing subcellular localization of numerous RNAs and proteins have been generally qualitative lacking detailed quantitative approaches to systematically describe the positions of RNAs and proteins. They have typically been constrained to a limited number of systems, in which spatial heterogeneity is controlled and subcellular partitions are easily defined. In developmental models, such as the *Drosophila* embryo and *Xenopus* oocyte, numerous mRNAs have been shown to localize to specific subcellular positions, which determine morphogen gradients and specify cell fates (Macdonald and Struhl, 1988, Tautz and Pfeifle, 1989). Similarly, a large number of mRNAs have been shown to be enriched in dendrites and synapses in neuronal systems, contributing to neuronal growth and establishing synaptic plasticity (Batish et.al., 2012, Buxbaum et al., 2014, Tzingounis and Nicoll, 2006). mRNA localization has also been shown in polarized cells of various kinds, such as budding yeast, and migrating fibroblasts (Martin and Ephrussi 2009, Mili et al., 2008), although typically only to coarsely defined regions, such as the leading edge of the cell. This is due to the diverse morphologies of such systems, as well as lack of methods to measure accurately specific subcellular domains.

Many of these model systems in addition to heterogeneity at the morphological level, lack quantitative approaches to spatially resolved omics and confront the problem of pervasive stochasticity at the level of gene expression (Raj et. al., 2006). A number of studies suggest that some of this stochasticity may be functional, for instance through the importance of higher-order distribution in grouping transcripts functionally (Battich et al., 2013), or as a compensating mechanism for differences in cell size (Padovan-Merhar et. al., 2015). Such functional stochasticity and noise thus add another level of complexity to developing quantitative analytical approaches to spatially resolved omics, since they need to capture such stochasticity explicitly. Although detailed mechanistic insights into RNA spatial and temporal positioning are emerging (Bouvrette et al., 2017, Moor et al., 2017, Zappulo et al., 2017, La Manno et al., 2018), a system that is able to capture and quantify dynamic RNA subcellular positioning, complementary to scRNA seq is required. Such a system would allow for a deeper understanding of cell states and assist in further identification and characterization of different cell types and sub populations. In sum, to unravel the mechanisms of RNA spatial and temporal distribution, quantitative tools that probe these relationships systematically need to be developed.

Here, we describe DypFISH, a spatially resolved omics approach overcoming the limitations above by quantitatively measuring the spatial distribution of slow-scale dynamics of mRNA and protein distributions at fine-grained spatial resolution in single cells. DypFISH leverages micropatterning which has been shown to lead to reproducible spatial organization of organelles (Schauer et al., 2010) and ensure that the cell size is known *a priori*, allowing the averaging of high number of cells. Thus this system deals with a number of significant sources of heterogeneity, which might interfere with the identification of spatial patterning and quantification. By selecting specific micropattern architectures, which mimic external constraints from a cellular environment, distinct patterns of subcellular localization of molecules of interest as well as spatial organization of organelles can be isolated and studied in detail (Théry et al., 2005, Théry et al., 2006). DypFISH builds upon the reproducibility of micropatterned cells to develop quantitative and testable models of RNA and positioning that can be extended to physiological contexts.

DypFISH introduces analytical techniques that allow joint analysis of discrete point-based single molecule Fluorescence In Situ Hybridization (FISH) mRNA data and continuous intensity immunofluorescence (IF) protein data. The analytical techniques include a generalized approach to identifying clustering dynamics, an approach to identifying dependencies of mRNA and protein spatial distributions on organelle positioning and an approach to identifying interdependent dynamics between mRNA transcripts and their corresponding protein products globally and at specific subcellular locations. Implementing DypFISH we uncovered fine-grained and reproducible aspects of localization dynamics, pointing to novel biological phenomena while revealing dynamic subcellular repositioning of RNA and proteins. DypFISH probes the dependencies uncovered through perturbation studies, thus allowing one to test for possible mechanisms underlying the localization dynamics and by extension changes in cell state, which we demonstrate in this study. Although we focused here on mRNA-protein subcellular localization, our approach is broadly applicable to spatially resolved omics of other molecular species, and scalable to incorporate high bandwidth such as MerFISH (Moffitt and Zhuang, 2016) where simultaneous assaying of subcellular localization of dozens of transcripts can be interrogated in a single cell.

## RESULTS

### Micropatterning of cells enhances reproducibility of mRNA subcellular distributions

We were particularly interested in the ability to interrogate subcellular positioning of RNA and protein in the context of altered cellular states. Therefore, we selected an established fibroblast system that allows the investigation of RNA positioning in relation to its polarity state (Mili et al., 2008). As mRNA candidates we selected a subset of mRNAs within a group of RNAs that had previously been identified as enriched in lamellipodia of fibroblasts upon polarization and cell migration (Hengst et al., 2009, Mili et al., 2008, Schmoranzer et al., 2009, Mili et al., 2008).

Micropatterning has been shown to lead to stereotypical localization of organelles, such as the centrosome, early endosomes, lysosomes and the Golgi apparatus (Schauer et al., 2010, Thery et al., 2005, Thery et al., 2006). We were interested in establishing whether micropatterning can similarly reduce variation in mRNA spatial distributions and enable us to construct a quantitative framework for measuring reproducible subcellular spatial localization of RNA and protein. To this end cells were induced to polarize on micropatterns and fixed at different time points post induction. We grew mouse fibroblasts on crossbow shaped micropatterns, which have been shown to be suitable for the study of polarizing cells (Schauer et al., 2010, Théry et al., 2006). Each slide was custom microfabricated to contain multiple 12 by 12 grids of crossbow-shaped micropatterns to which the cells adhered (Figure S1A). We developed an autonomous image acquisition and semi-automated image analysis pipeline (Figure 1A) able to scan each microfabricated slide and autonomously acquire images of individual cells at high magnification. We used either standard wide-field fluorescent microscopy or spinning disk confocal microscopy at to acquire a three-dimensional stack of images for each cell.

**Figure 1.**
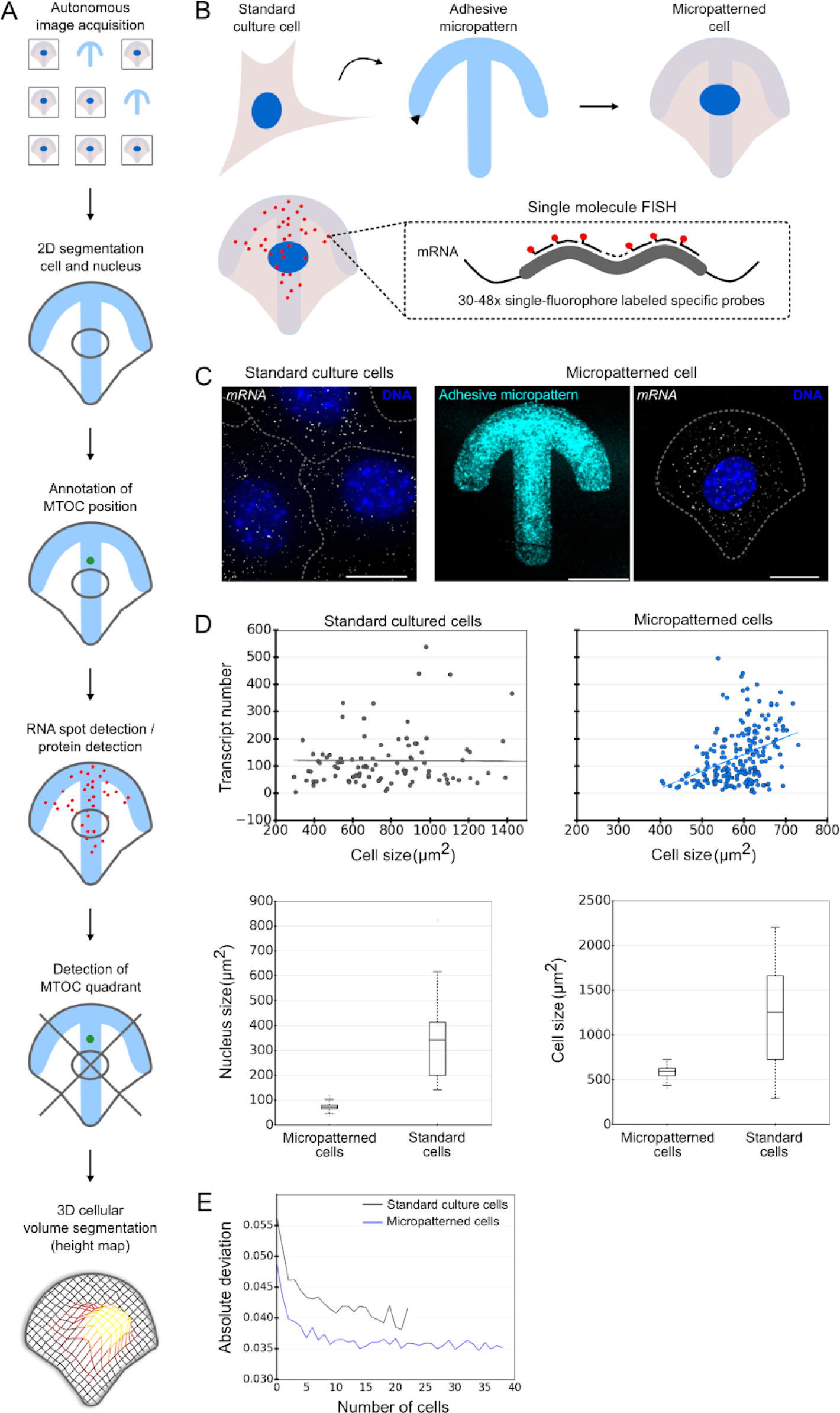
Reproducibility of mRNA and protein distributions in micropatterned cells. (A) Mouse fibroblasts were plated on fibronectin-coated micropatterns and induced to polarize by addition of serum. Cells were fixed and single molecule FISH was performed to target mRNAs of interest. (B) Outline of the image processing pipeline. *Arhgdia* mRNA was visualized using single molecule FISH (grey) in standard and crossbow-shaped micropatterned mouse fibroblasts, micropatterns were visualized by coating with labeled fibrinogen (cyan) and DNA was stained with Hoechst (blue), scale bar 10 μM. (D) Comparison of the relationship between cell size and Arhgdia transcript copy number in standard cultured and micropatterned cells. Solid lines in upper graphs show the least squares fit. Lower graphs compare cell and nucleus size in the two conditions. (E) Absolute deviation of Arhgdia mRNA distribution of a randomly selected cell from a pooled average of up to ~40 cells for cultured and micropatterned cells.

Single molecule FISH (Raj et al., 2008) and IF were performed to label mRNAs and corresponding proteins of interest respectively (Figure 1B). We also labeled the microtubule (MT) cytoskeleton, the nucleus and micropattern base. Representative images of the micropatterned base and single molecule FISH are shown in Figure 1C. A complete list of the acquired images is shown in Supplementary Table 1. To extract information of interest from the microscopy images, we built a custom image analysis pipeline with (1) manual annotation of the MTOC position, (2) 2D automated segmentation of the cell and nucleus areas, (3) automated spot detection and (4) height-map construction across a stack of 2D cellular regions to define a 3D segmentation of the cell volume (Figure 1A; Experimental Procedures for computational details for steps 3, 4 and 5).

To investigate the localization dynamics of mRNAs and proteins translated from these transcripts we followed specific mRNAs and their respective proteins at different time points (2, 3, 4 and 5 hours for mRNA, and 2, 3, 5 and 7 hours for protein) post induction of polarization by serum. These time points were chosen as fibroblasts polarize over this time scale on crossbow micropatterns. In order to determine whether micropatterning leads to reduced heterogeneity in mRNA transcript and protein distributions, we compared fibroblasts grown in standard culture with those grown on micropatterns using the same experimental pipeline, analyzing only cells lying fully in the field of view. Representative images of standard cultured and micropatterned cells are shown for the *Arhgdia* mRNA, which was previously found to be enriched at the leading edge of polarized fibroblasts (Mili et al., 2008) (Figure 1C).

First, we compared the reproducibility of *Arhgdia* mRNA distributions in standard cultured and micropatterned cells by spatially quantizing the transcript distributions across a grid of regular voxels. The absolute deviation of the quantized distribution of a randomly selected cell from a pooled average is reduced in micropatterned cells for all pool sizes up to ~40 cells (Figure 1E). The error profiles of these distribution descriptors for this transcript are concordant with a previous study, which estimated that ~20 micropatterned cells were necessary to establish reproducible organelle positions using the AMISE metric (Schauer et al., 2010). We further investigated the impact of micropatterning on the volume-corrected noise measure *Nm* introduced in a previous study (Padovan-Merhar et al., 2015). Consistent with that study, we observed linear relationships between transcript number and cell size in both standard cultured and micropatterned cells, as demonstrated for *Arhgdia* mRNA (Figure 1D). However, the linear relationship is less stochastic in the micropatterned cells, leading to a lower *Nm* value (Figure 1D). Further comparison revealed a tighter distribution of cell and nuclear sizes in the micropatterned cells, consistent with a mechanism for cell size determination, which relies on low variability of nuclear size, as proposed previously (Figure 1D) (Padovan-Merhar et al., 2015).

We further investigated the profiles over time of the *Nm* for a series of mRNA transcripts including *Pkp4* and *Rab13*, which are enriched at the leading edge in polarized fibroblasts (Mili et al., 2008), *Pard3*, which translates into the Par3 protein that controls different aspects of polarity in various cell types and is enriched in developing axons (Hengst et al., 2009, Schmoranzer et al., 2009), *β-Actin*, a well studied localized mRNA in various cell types and *Gapdh*, which to the best of our knowledge is not known to localize to specific subcellular domains. We found for a number of transcripts a reduction in *Nm* over time, up to the 4 h time point. (Figure S1C). These data strongly indicated that micropatterns conveyed important advantages amenable to quantitative analysis of RNA position over standard cell culture.

### A subset of mRNAs and corresponding proteins shows peripheral enrichment and correlated clustering dynamics

We investigated the joint localization dynamics for mRNA and corresponding protein associated with four of the mRNA transcripts above (*Arhgdia*, *Pard3*, *β-Actin* and *Gapdh*) and the mRNA localization dynamics for the remaining two transcripts (*Pkp4* and *Rab13*). We developed a cell quantization method for local enrichment statistics as well as a temporal interaction score to measure the interdependence between mRNA and protein dynamics. Initially, we characterized whether the mRNA and corresponding proteins are enriched in the periphery of the cells by calculating the fraction of cytoplasmic transcripts, which lie within a band at the boundary of the cell, whose width is a fixed proportion of the radial distance to the nucleus edge (Experimental Procedures). A subset of the transcripts, including *Arhgdia* and *Pkp4*, which were previously shown to be enriched at the leading edge (Figure 2A) (Mili et al., 2008), as well as *Pard3*, whose localization in fibroblasts has not been characterized previously, are peripherally enriched for up to 30% of the radial distance (Figure 2B). Gapdh mRNA and protein characteristic distributions were used as a control as neither the mRNA nor protein were expected to show patterns of enrichment or strongly localized dynamics (Mili et al., 2008).

**Figure 2.**
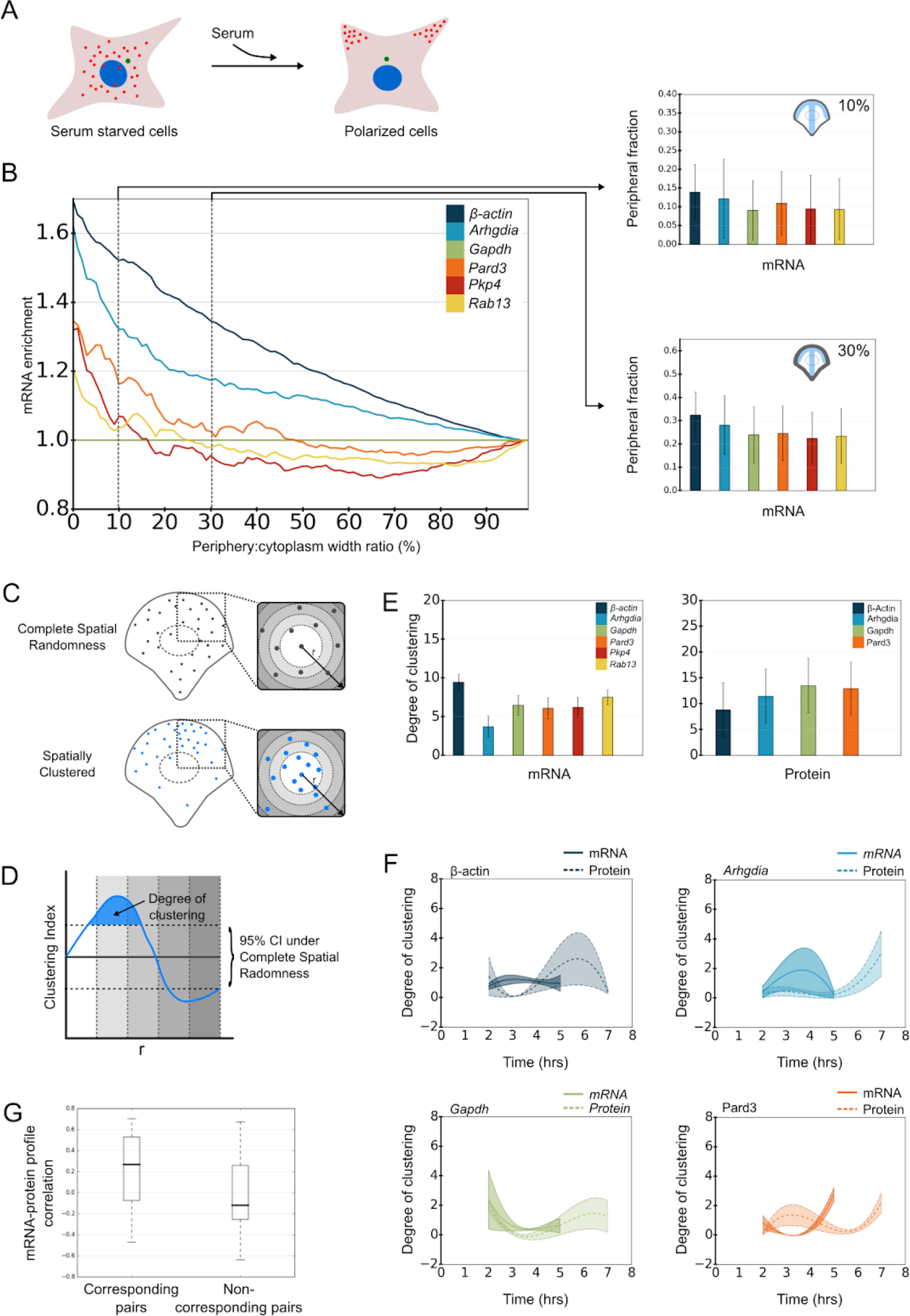
Peripheral enrichment and clustering dynamics of mRNA-protein pairs. (A) A subset of transcripts were previously shown to be enriched in the leading edge of mouse fibroblasts upon treatment with LPA/serum (Mili *et al.*, 2008). (B) Comparison of the enrichment of 5 mRNAs with respect to the *Gapdh* mRNA in a peripheral cellular region whose width varies from 0-100% of the radial distance from the plasma membrane to the nucleus (left), and the distributions of absolute fractional values at 10% and 30% (right). (C) Clustering is characterized by comparing observed transcript and protein distributions to complete spatial randomness. For mRNAs, the Ripley’s K function is estimated for an observed distribution and samples from a homogeneous Poisson process by counting the number of pairs of points lying within a radius *r* of each event. (D) To compute the degree of clustering, we first defined an estimator of the Ripley’s K function, called clustering index, by normalizing the observed Ripley’s K function to the 95th and 5th percentiles under the Poisson process. Statistically significant clustering of mRNAs or proteins is found at radius r where the estimator function is over the 95th percent confidence interval calculated based on the CSR assumption. Values below the 95th percent confidence interval indicate the dispersion of the molecules. The degree of clustering is the area under the estimator’s curve that is above the 95th percentile of the random distribution. (E) Comparison of degree of clustering for mRNAs and proteins (all time-points, log values shown after scaling by log(0.5) and log(0.01) for mRNAs and proteins respectively). (F) Comparison of clustering dynamics for four mRNA-protein pairs using degree of clustering. Zero values indicate distribution at a given time-point is not distinguishable from randomness. (G) Correlations between temporal profiles for corresponding and non-corresponding mRNA-protein pairs (Pearson Correlation Coefficient used). Correlations are between median values at 2, 3 and 5 h time points for degree of clustering, peripheral fraction, cytoplasmic total and spread descriptors. Bar graphs in (B) and (E) show median with .25 and .75 quantile error bars, and graphs in (F) show median surrounded by envelope indicating .25 and .75 quantiles fitted to cubic splines. See also Figure S2.

We next analyzed the clustering behaviour of mRNAs and proteins using a generalization of the Ripley’s K analysis (Lee et al., 2013, Ripley, 1977) that we first introduced in (Warrell et al., 2016). Ripley’s K is a commonly used algorithm to describe the extent of clustering of points, such as mRNAs (Figure 2C). For each mRNA spot and each distance *d* the number of transcripts lying within a sphere of radius *d* is counted. Spatial clustering of mRNAs can then be calculated by estimating the probability distribution of this function under a null hypothesis of complete spatial randomness (CSR) and comparing it with the function calculated from observed (spatially clustered) transcripts. We adjusted (Experimental Procedures) the algorithm based on the generalized Ripley’s K function for evaluating the extent of clustering of both mRNA (discrete) and protein (continuous) spatial distributions. This was done by computing the *degree of clustering*, a unitless measure, that can be used to compare clustering between different molecules and conditions. We summed the area where the normalized Ripley’s K function deviates from the 95 % confidence interval of the random distribution (Figure 2D).

We evaluated the degree of clustering of all transcripts and proteins across all time points, revealing high overall values for all proteins and various values for the different mRNA transcripts (Figure 2E). We further calculated the degree of clustering at each individual time-point for *Arhgdia*, *Gapdh*, *β-Actin* and *Pard3* mRNAs and corresponding proteins. By visualizing the mRNA and protein profiles, a relationship is suggested for the Arhgdia, β-Actin and Pard3 mRNAs and proteins, which show a peak in the mRNA profile followed by a peak in the proteins, strongly suggesting temporal causality (Figure 2F).

We then calculated additional basic mRNA/protein distribution descriptors such as cytoplasmic total transcripts/intensity, peripheral fraction and cytoplasmic spread (Figure S2A, and Experimental Procedures). We compared the mRNA and protein profiles of these descriptors for all time points and found that many of these show related values for the same gene (corresponding pairs) (Figure S2B). The observed differences between the comparisons of corresponding and non-corresponding mRNA-protein pairs across all descriptors are shown in Figure 2G and are statistically significant. The analysis of basic distribution descriptors as well as clustering dynamics, for several of the mRNA-protein pairs, suggests they exhibit interdependent spatial positioning. Using the tools discussed above, we were thus able to extract significant subcellular spatial information for different mRNA and protein species.

### Localization of a subset of mRNAs and corresponding proteins shows strong correlative influence of MTOC position

Several components of cellular architecture are known to change dynamically during polarization, including the cytoskeleton and the microtubule organizing center (MTOC). The MTOC marks the center of most eukaryotic cells and is associated with the position of the nucleus. In the majority of polarized cell systems and developing neurons, the MTOC is positioned between the nucleus and the leading edge prior to migration or local cell growth (Gomes et al., 2005, Hale et al., 2011). We reasoned that if cells were to reliably and reproducibly position RNA or protein within the subcellular volume, then this process could be linked to an ability to sense the MTOC position. Whether mRNA and its corresponding proteins are subject to such reorientation is unknown. Having demonstrated that several mRNA transcripts and proteins exhibit significant and reproducible clustering dynamics, we sought to relate this behavior to key cellular structures and organelles. Initially we were interested in probing the subcellular spatial position of RNA and protein relative to the position of the MTOC. More specifically, we sought to determine whether the clustering dynamics we observed were dependent on MTOC positioning. For our analysis, we divided the cell into quadrants, which we used as regions over which we could estimate mRNA and protein local densities (Figure 3A) and we also determined the MTOC localization within these quadrants (Figure 3B). We observed higher enrichment of all cytoplasmic mRNA transcripts in the MTOC-containing quadrant (Figure 3C). Additionally, *β-Actin*, *Pard3*, *Pkp4* and *Rab13* transcripts were specifically enriched in the MTOC-containing quadrant when located in the leading edge of the cell. Similarly, all peripheral transcripts showed higher enrichment in the MTOC-containing quadrant, with this enrichment being more distinct compared to the cytoplasmic population. *β-Actin*, *Gapdh*, *Pkp4* and *Rab13* showed clear enrichment in the MTOC-containing quadrant when positioned in the leading edge of the cell.

**Figure 3.**
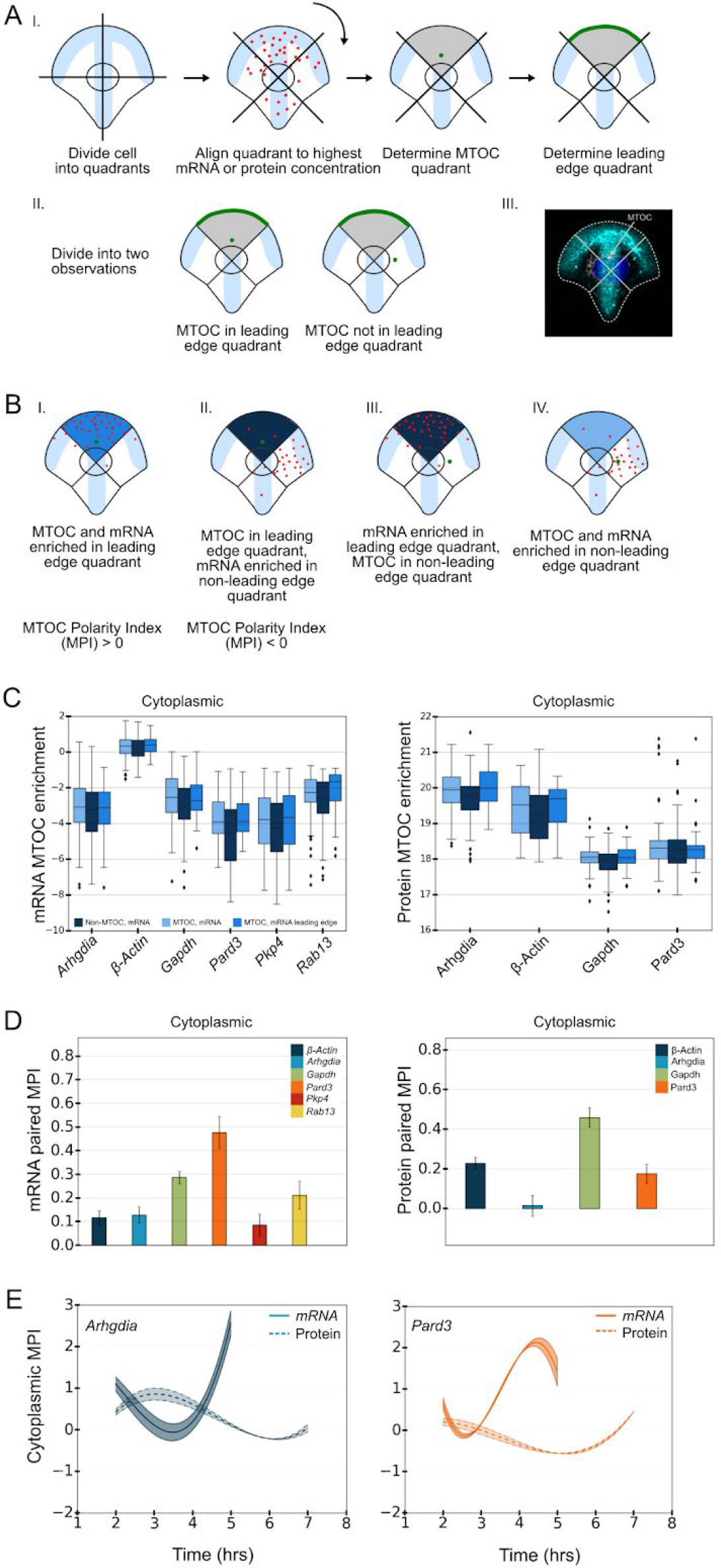
Correlative influence of cytoplasmic mRNA and protein distributions and MTOC position. (A) Schematic of the MTOC correlative influence analysis. MTOC position is annotated in projected 3D Tubulin IF images (iii). (B) Schematic of the analysis to determine the MPI value. The MTOC polarity index (MPI) is defined by normalizing the differences of signal concentration between the MTOC associated quadrant and the other quadrants. It takes values between −1 and +1, with positive values indicating enriched mRNA or protein concentration in the MTOC quadrant, negative values indicating enrichment away from the MTOC quadrant, and values close to 0 indicating no correlative enrichment. (C) The cytoplasmic mRNA enrichment in non-MTOC containing quadrants, MTOC-containing quadrants and MTOC-containing quadrants when this quadrant coincides with the leading edge. The concentration of cytoplasmic *β-Actin*, *Pad3*, *Pkp4* and *Rab13* transcripts and Arhgdia and β-Actin proteins is enriched in the MTOC-containing quadrant when it is in the leading edge. (D) Comparison of MPI values for mRNAs and proteins in cytoplasmic populations (all time points). (E) Comparison of MPI dynamics for Arhgdia and Pard3 mRNA-protein pairs. Bar graphs in (D) show median and .25 and .75 quantile error bars for 11 bootstrapped MPI estimates. Graphs in (E) show median surrounded by envelope indicating .25 and .75 quantiles of 11 bootstrapped estimates fitted to cubic splines.

Based on the above observations, we were able to introduce an *MTOC Polarity Index* (MPI) to analyse MTOC dependent enrichment in both mRNA and protein distributions. This indicator lying between −1 and +1 is derived by normalizing the differences of signal concentration between the MTOC associated quadrant and the other quadrants. Positive MPI values imply MTOC dependent enrichment of RNA transcripts, negative values imply enrichment away from the MTOC, and a value of zero implies no detectable correlative influence (Figure 3D). The MPI can be calculated for both point based and continuous valued measurements, corresponding to mRNA and protein distributions respectively, and significant enrichment can be identified by a statistical test against CSR (Experimental Procedures).

We calculated the MPI as above for all transcripts (Figure 3D) and proteins (Figure S3D), calculating values for both the whole cytoplasmic population, and the peripheral population at 10 % radial distance, using all time points. *Pard3, Rab13* and *Gapdh* mRNAs show significant MPI scores in the cytoplasmic population, whereas all mRNAs, excluding *Gapdh* show significant MPI scores in the peripheral population, suggesting that the MTOC orientation affects the localization of these mRNAs as it changes during polarization (Figure 3D). Finally, we calculated MPI scores across time for all mRNA-protein pairs (Figure 3E and Figure S3E) and observed profiles suggesting a temporal correlative influence between mRNA, protein and MTOC orientation for Arhgdia and Pard3.

### Dynamics of mRNA-protein distributions are consistent with MTOC-dependent patterns of localized translation

As previously described, several mRNA-protein pairs appeared to show interdependent distributions for the basic descriptors, clustering indices and MPI. The interdependency could reflect spatially and temporally restricted translation (local translation) (Besse and Ephrussi, 2008) and/or separate localization pathways for mRNAs and proteins to common subcellular locations. To explore these interdependencies we derived a *Temporal Interaction Score* (TIS), which is a value between 0 and 1, computed as the normalized rank-sum of the correlations between mRNA and later protein distribution pairs in a ranking across all pairs of time-points (See for details in Experimental Procedures III.7). A large TIS value for a pair of corresponding mRNAs and protein is consistent with interdependent dynamics, and may suggest local translation, although it does not rule out alternative mechanisms such as separate mRNA and protein localization pathways with delayed protein transport.

A TIS can be calculated for any measure of correlation between mRNA and protein distributions, which allowed us to probe for interdependent dynamics with respect to specifically defined subcellular regions. We spatially quantized the cells (i) radially with the center at the nucleus centroid and (ii) circularly by computing isolines at different distances from the cell’s periphery (see Experimental Procedures section III.7 and Figure 4A(i)). We thus obtained a fine grained quantization of each cell into segments, and were able to compute subcellular spatial distribution profiles of mRNAs and proteins, corresponding to concentration statistics in each segment (Figure 4A(ii)). The latter analysis was prompted by the observation that our peripheral MPI scores are relatively stronger for proteins than mRNAs. While both showed strong cytoplasmic MPI scores (Figure 3D), we may particularly expect interdependencies between cytoplasmic mRNAs and peripheral and/or cytoplasmic proteins in a common direction with respect to the MTOC. This could potentially reflect radial mRNA transport on the cytoskeleton in preferred directions leading to protein enrichment in those directions at the periphery due to local translation. We computed TIS values using (a) *global* correlations of all voxels/segments across the cytoplasmic area, and (b) *local* correlations across subsets of voxels/segments within peripheral regions (Figure 4C). We then calculated global TIS values for four corresponding mRNA-protein pairs (Arhgdia, Gapdh, β-Actin and Pard3) using for comparisons the ‘forward-leading’ time point pairs shown in (Figure 4B). Using the fine grained quantization scheme, we observed significant interdependent dynamics for all cytoplasmic mRNA-protein pairs (Figure 4D).

**Figure 4.**
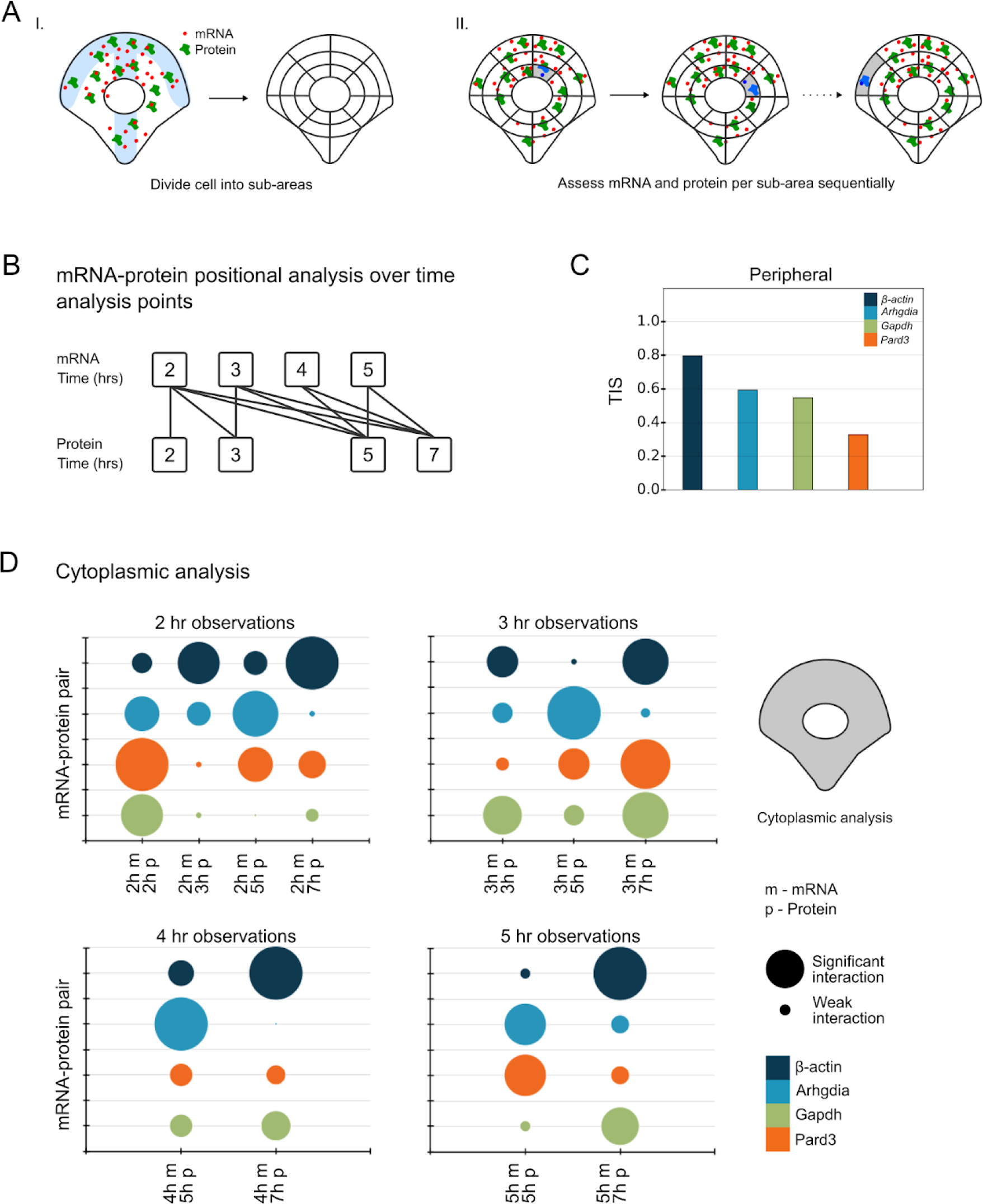
Interdependency of localization dynamics for corresponding mRNAs and proteins. (A) The Temporal Interaction Score (TIS) as a correlation between mRNA and protein spatial distributions for image acquisitions at several time points. Distributions were spatially quantized radially with the center at the nucleus centroid and circularly by computing isolines at different distances from the cell’s periphery. The subcellular distribution profiles of mRNA and proteins corresponding to concentration statistics in each segment was computed. (B) Forward leading time point pairs, defined as *t*_1_ < *t*_2_, were used for calculating TIS values. (C) TIS values were computed using global correlations of all voxels/segments across the cytoplasmic area, and local correlations across subsets of voxels/segments within peripheral regions. (D) Significant interdependent dynamics for all cytoplasmic mRNA-protein pairs was observed using the fine grained quantization scheme.

### Perturbation of various cytoskeletal components disrupts characteristic mRNA-protein localization and interdependency patterns and hints at local translation

We were generally interested in quantitatively measuring cell state by using mRNA copy number and incorporating dynamic changes to subcellular positions over time. In particular we wanted to include a measurement of mRNA-protein dynamics that could reveal interdependencies influencing their subcellular positions.

Our analysis had already revealed interdependent mRNA-protein dynamics, which suggested local translation. However, as we could not rule out independent localization of mRNA and corresponding proteins, we introduced two perturbations to inhibit potential transport pathways of the different molecular species. First, we disrupted microtubule polymerization using nocodazole (Figure 5A), which we reasoned would lead to disruption of the MTOC dependencies of selected mRNA and proteins, as well as potentially loss of local translation. We selected Arhgdia and Pard3 mRNA-protein pairs to test this hypothesis and collected mRNA FISH and protein IF data at 3 hours and 5 hours post exposure to nocodazole and compared it with untreated data as above from the equivalent time points.

**Figure 5.**
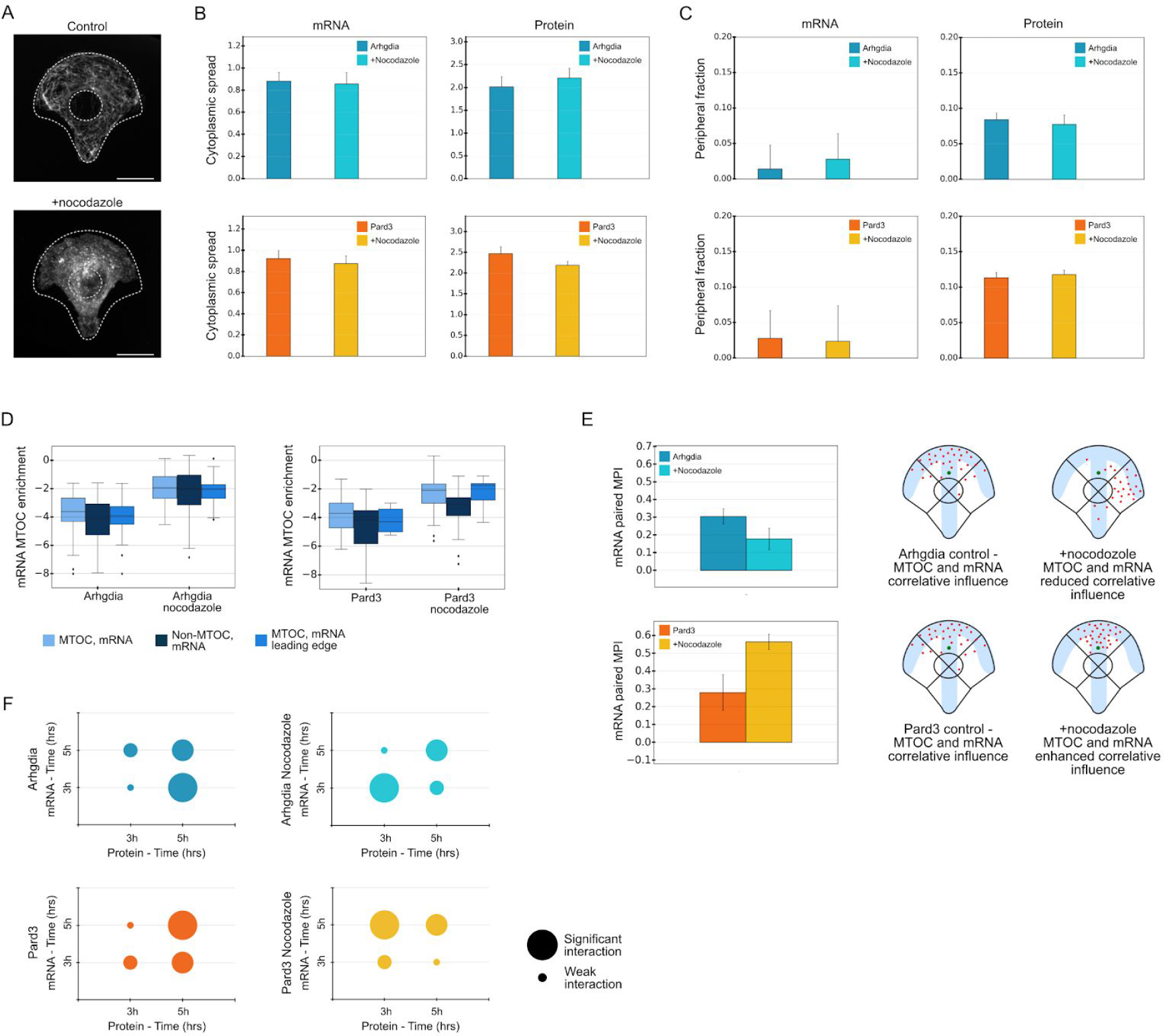
Effects of cytoskeleton disturbance on mRNA-protein localization and interdependent dynamics. (A) Nocodazole was added to cells seeded on micropatterns, inducing inhibition of microtubules polymerization (Tubulin IF image shown), scale bar 10 μM. (B) Cytoplasmic spread of Arhgdia and Pard3 transcripts and proteins (at 3 and 5 h combined) is defined as a statistics measuring the evenness of a molecule spread across the cell, with the value 1 for even distribution. No significant effects in cytoplasmic spread are observed for these two genes in the presence of nocodazole; (C) The peripheral fraction of Arhgdia and Pard3 transcripts and proteins (at 3 and 5 h combined) were calculated similarly to Figure 2B. *Arhgdia* peripheral fraction was increased in the presence of nocodazole. (D) mRNA MTOC enrichment profiles for *Arhgdia* and *Pard3* shown for nocodazole-treated and control cells at 3 and 5 h combined. A reduction in MTOC enrichment for *Arhgdia* and an increase in MTOC enrichment for *Pard3* are observed in the presence of nocodazole. MPI scores shown for control and nocodazole treated cells for *Arhgdia* and *Pard3* transcripts at 3 and 5 h combined. Nocodazole treatment disrupts characteristic dependency between MTOC orientation and localization, resulting in lower MPI values than in the untreated conditions for *Arhgdia* and higher MPI values compared to untreated conditions for *Pard3*. (F) The interdependent dynamics for Arhgdia and Par3 were disrupted in the presence of nocodazole.

We first calculated the effects of nocodazole treatment on the total mRNA count (Supplementary Figure 3A), the cytoplasmic spread of the transcript and the corresponding protein (Figure 5B) and the 10 % peripheral fraction (Figure 5C). *Arhgdia* cytoplasmic spread was not significantly altered upon treatment with nocodazole, whereas its peripheral fraction was decreased (Figure 5B and 5C). RhoGDI’s (the protein translated from *Arhgdia*, to which, we refer as Arhgdia for simplicity) cytoplasmic spread was slightly increased whereas the peripheral fraction of the protein was not significantly affected (Figure 5B and 5C), suggesting that Arhgdia mRNA and protein do not make exclusive use of the microtubule network for transport. The cytoplasmic spread and peripheral fraction of *Pard3* were not significantly changed upon treatment with nocodazole (Figure 5B and 5C). A slight reduction was detected in the cytoplasmic spread of Pard3 protein in the presence of nocodazole (Figure 5B) and no significant change in the peripheral fraction of the protein (Figure 5C), suggesting that similarly to *Arhgdia*, *Pard3* and its corresponding protein do not make exclusive use of the microtubule network.

Next, we probed the effects of the nocodazole treatment on the correlative influence of the polarization of the cell and the subcellular localization of the mRNA transcripts and their corresponding proteins (Figure 5D and 5E). The MTOC enrichment and MPI of *Arhgdia* in nocodazole-treated cells compared with control cells were significantly reduced (Figure 5D and 5E), indicating that the orientation of the mRNA subcellular localization was lost. This suggestested that though the prior calculation of cytoplasmic spread showed only modest effects of the drug, the influence of the MTOC had become decoupled from the subcellular positioning of *Arhgdia* RNA. Strikingly, the opposite effect was observed for *Pard3*, where a significantly higher MTOC enrichment and MPI were observed for the mRNA in the nocodazole treated cells (Figure 5D and 5E), indicating that the perturbation to the MTOC had the effect of tightening the influence of MTOC position on RNA distribution.

We then probed the effects on mRNA-protein interdependency by calculating local TIS maps for control and nocodazole treated cells on both mRNA-protein pairs, restricted to the 3-5 hours time points (Figure 5F). The interdependency for Arhgdia mRNA-protein pair was disturbed, possibly stemming from the decoupling of the MTOC influence on the subcellular distribution of the RNA and protein in nocodazole-treated cells. In contrast, nocodazole did not have a significant effect on the interdependency for the Pard3 mRNA-protein pair, which could correlate with a higher correlative influence of the MTOC on *Pard3* mRNA localization observed in nocodazole-treated cells.

Other cytoskeletal networks could influence RNA subcellular positioning. To probe these, we next treated the cells with the actin polymerization inhibitor cytochalasin D (cytoD) for 1 hour before fixation of cells and compared FISH and IF data for the Arhgdia mRNA-protein pair. Similarly to the treatment with nocodazole, *Arhgdia* mRNA’s cytoplasmic spread remained unaffected (Supplementary Figure 3B), whereas the concentration of the transcript was higher on the periphery of cytoD-treated cells as compared to control cells (Supplementary Figure 3C). Similarly to the mRNA, the cytoplasmic spread of Arhgdia was not significantly affected in the presence of cytoD (Supplementary Figure 3B) however, the peripheral concentration of the protein was decreased in the cytoD-treated cells (Supplementary Figure 3C). These data suggest that either local translation at the periphery of *Arhgdia* was affected by the drug, or that mechanisms anchoring Arhgdia to the periphery are lost in the presence of the drug.

### Sarcomeric mRNAs cluster in a striated pattern in differentiated myofibers

In order to further validate Dyp-FISH, we sought a cellular model where RNA subcellular localization and potential local protein translation linked to dynamic changes in cells state could be interrogated. We focused on the unusually large multinucleated muscle cells termed myofibers. These large cells with tubular shape are formed by fusion of mononucleated cells and their main function is to generate mechanical force via contraction. Muscle contraction is achieved by the shortening of sarcomeres that are organized along the length of the myofiber. Each sarcomere is flanked by a Z-line, the site of anchoring of the actin filaments, thus resulting in the striation of the myofiber (Franzini-Armstrong and Peachey, 1981). We used skeletal muscle since the myofiber has an invariable tubular shape and a highly predictable cytoplasmic organization. The myonuclei in these cells are spaced at regular intervals, an important feature for muscle function (Bruusgaard et al., 2003, Bruusgard et al., 2006; Manhar, 2018). *In vitro* differentiation of myofibers allows for high resolution imaging throughout distinct developmental stages, including the formation of patterned sarcomeres with well defined z-line striations (Falcone et al., 2014; Pimentel et al., 2017; Vilmont et al., 2016) permitting the capture of dynamic changes in cell state reflected in RNA-protein subcellular localization. Thus, the size and regularity of myofiber architecture allied with a temporal component make them an excellent candidate to compute spatial distribution profiles of mRNAs and proteins.

With this aim, we analyzed the distribution of an mRNA that encodes a protein found at the z-lines, during muscle differentiation. We choose the *actn2* mRNA, which encodes for α-actinin, the main component of z-lines relative to the sarcomeric Z-lines (Figure 6A). To our surprise, the majority of the *actn2* mRNA was found in the vicinity of the Z-line in mature myofibers (Figure 6B). In order to understand if this clustering depends on the developmental stage of the myofiber we imaged immature myofibers in which the Z-lines are less organized. Additionally, we also probed the *gapdh* mRNA distribution as a non-sarcomeric control in both immature and mature myofibers. The degree of mRNA proximity was lower in both cases, suggesting that possibly *actn2* mRNA localization precedes protein organization (Figure 6B) of the Z-line. To better address this question we quantized the images perpendicularly to cell axis, similarly to how sarcomeres are organized (Figure 6C). The highest degree of clustering was observed for *actn2* mRNA in mature myofibers, when compared to the immature counterpart or to the *gapdh* mRNA. These quantitative data strongly suggest that the *actn2* mRNA distribution specifically follows the respective protein organization, instead of preceding it (Figure 6D). These data shed light on a long standing question in the field and produce a basis of testable hypotheses for how *actn2* mRNA is directed to the Z-line.

**Figure 6.**
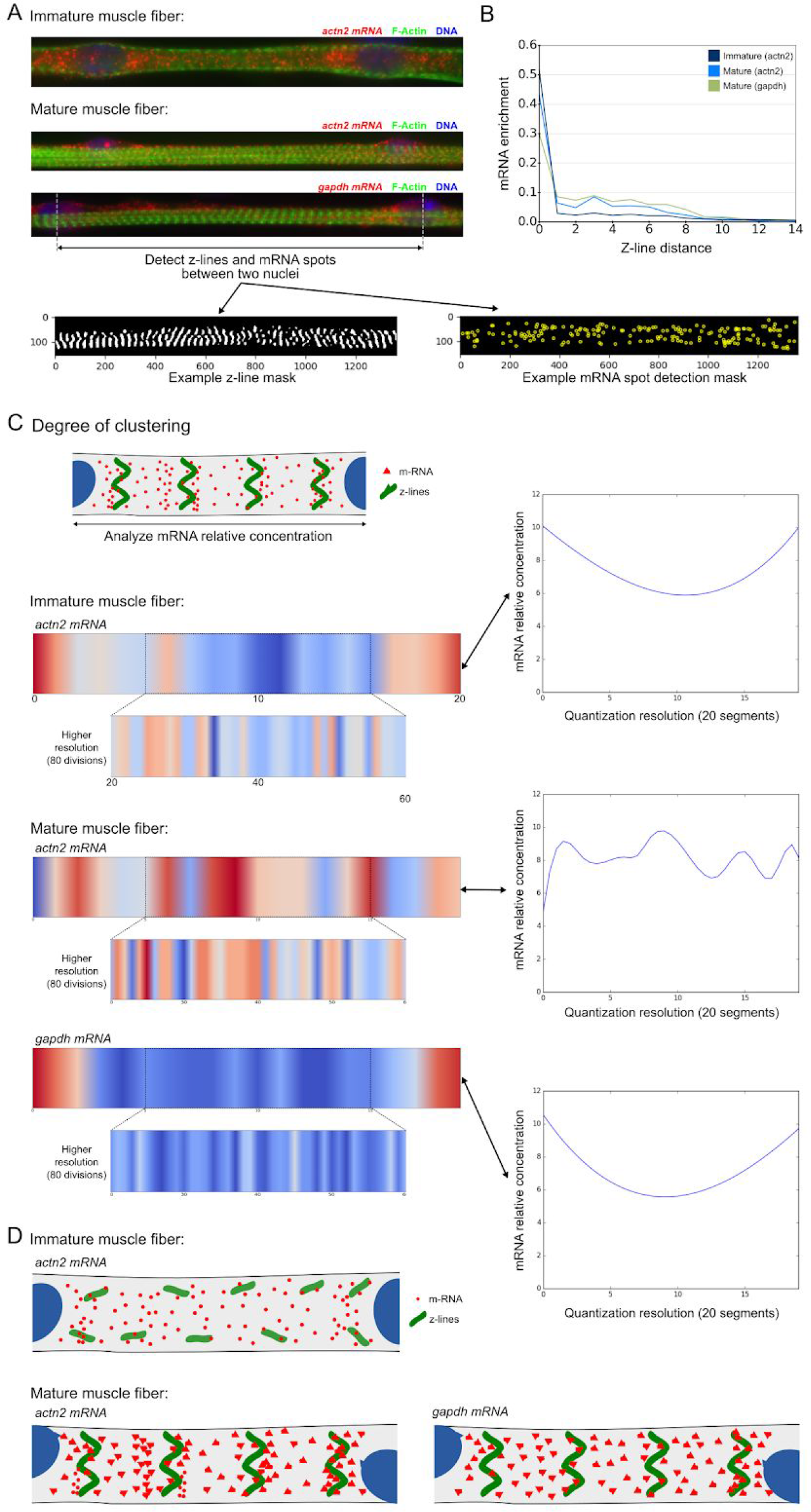
Sarcomeric mRNAs cluster in a striated pattern in differentiated myofibers. (A) Typical epifluorescent images of immature and mature muscle fibers. The DNA was stained with DAPI (blue), F-actin was visualized using immunofluorescence (green) and *Actn2/Gapdh* were visualized using single molecule FISH (red). Z-line and RNA spot detection masks were extracted using immunofluorescence and single molecule data respectively. (B) mRNA distance profiles. For each mRNA we computed its distance to the closest Z-lines, which allowed us to count the number of mRNAs having a certain distance to Z-lines. Normalized median counts are represented on the y axis. A higher number of *actn2* immature *mR*NA falls inside or close to Z-lines compared to mature fibers, suggesting greater clustering of mRNA between Z-lines for mature *actn2*. (C) The mRNA local density was computed between two nuclei. Each cell was quantized in vertical quadrants and relative concentration of mRNA in each quadrant was computed by normalizing the counts by the relevant surface. A wave-like clustering is observed for *actn2* in mature compared to immature fibers. No clustering is observed for *Gapdh*. (D) Model describing *actn2* mRNA distribution in immature and mature fibers.

## DISCUSSION

A wide variety of methods have been proposed for studying RNA localization with subcellular accuracy, including microscopy-based methods, such as those based on FISH (Battich et al., 2013, Chen et al., 2015, Lecuyer et al., 2007) or padlock probes (Larsson et al. 2010), as well as non-microscopy based methods such as Transcriptome In Vivo Analysis (TIVA) (Lovatt et al. 2014) and RNA Tomography (Junker et al. 2014). While there has been much interest in identifying patterns of localization from such data, a number of factors limit the generality of the kinds of analysis employed. Although developmental systems can be straightforwardly aligned in space and compared across time for comparison of expression patterns (Junker et al. 2014, Lecuyer et al., 2007), this is not always the case in other systems. In systems with greater heterogeneity, approaches to identify RNA spatial patterning have not generally attempted to take polarization or organelle arrangement into account (Battich et al., 2013, Chen et al 2015) and have not considered temporal patterning. Further, some of the most comprehensive approaches have relied on manual annotation of visual features to search for correlations between RNA patterns and RNA-protein patterns (Lecuyer et al., 2007) and hence lack a principled approach for the identification of correlations that may not be visually apparent. We set out to develop a quantitative method for investigating the spatial and temporal distribution of RNA and protein. To achieve this, we hypothesized that the use of micropatterning would reduce cellular heterogeneity and enhance the reproducibility of spatial distributions. We were able to achieve this in combination with automated high content imaging based on RNA FISH and IF labelling, and a range of specialized analytic techniques. We have used our approach to interrogate mRNA and protein spatial and temporal distribution in polarized fibroblasts and elucidate the methods by which particular mRNAs and proteins are localized, revealing a general dependence of mRNA-protein localization and dynamics on MTOC orientation. Through perturbation studies, we have demonstrated DypFISH’s ability to quantitatively detect changes in localization behaviour, confirming both the robustness of the approach and its ability to test mechanistic hypotheses.

### Shared RNA subcellular distribution patterns and the deterministic role of the MTOC

The general dependence of different mRNA species on MTOC orientation and the similarities in localization and dynamics of those associated with the leading edge, indicate specific subcellular distribution patterns in accordance with broader processes, which are also responsible for controlling nucleus and MTOC relative orientation during polarization (Kim et al., 2014, Razafsky et al., 2014). We posed the question whether proteins translated from mRNAs that consistently localize to similar subcellular locations are translated in these locations. By revealing the existence of mRNA-protein interdependencies, which are highly localized, many have a high correlative influence with the MTOC orientation and can be impaired by applying various perturbations, with our results pointing to local translation. In addition to the above, our results have general implications for stochasticity of gene expression in morphologically constrained biological contexts. The pronounced reduction in the variability of distribution descriptors we were able to achieve using micropatterning suggests that tissues such as gut epithelia, skin and others of constrained but similar morphology, may have far less transcriptional heterogeneity than previously thought.

### *Arhgdia* and *Pard3* localization

We were able to reveal significant aspects of the spatial and temporal distribution of specific mRNA transcripts and proteins of interest, such as *Arhgdia* and its protein product RhoGDI, a key factor in the Rho/Rac/Cdc42 (Rho GTPases) pathway. The transcript and protein show MTOC-dependent interdependency patterns, possibly indicating MTOC-dependent local translation. The RhoGDI protein is a negative regulator of the Rho GTPases, which are involved in a range of important cellular processes such as polarization, regulation of cytoskeletal organization, cell growth control and many others (Machacek et al., 2009, Sadok and Marshall, 2014, Schaefer et al., 2014, Zegers and Friedl, 2014). We showed here that the majority of the *Arhgdia* mRNA population is both cytoplasmic and MTOC-dependent, mainly located in the perinuclear area corresponding with the ER (data not shown), with a small fraction being localized to the periphery. This latter fraction may reflect a population that is transported to the periphery which can then be quickly translated when needed in order to regulate the levels of active Rho GTPases locally and in a rapid manner at the leading edge of the cell. The peripheral MTOC-dependent TIS maps for *Arhgdia*, as well as a decrease in the concentration of RhoGDI (Arhgdia) at the cell periphery despite minimal disruptions to the transport of both *Arhgdia* mRNA and RhoGDI protein suggest MTOC-dependent peripheral local translation, that may be required as part of the polarization process.

Our analyses also revealed important aspects of Pard3 spatial and temporal distribution. The localization of both transcript and protein show significant interdependent dynamics, implying that mRNA localization and localized translation may be factors which influence localization of the Pard3 protein during polarization. In contrast to *Arhgdia*, there is an increase in the MPI of *Pard3* mRNA upon disruption with nocodazole, indicating MTOC-MT influence on *Pard3* transport and peripheral anchoring of *Pard3*. Localization of the Pard3 protein in peripheral clusters at cell-cell adhesions was previously shown in fibroblasts when grown in culture (Schmoranzer et al., 2009), which is concordant with the peripheral enrichment of *Pard3* mRNA and protein in our system. Further, localization of the *Pard3* mRNA has been shown in growing axons stimulated with NGF and netrin-1 (Hengst et al., 2009). However, to our knowledge *Pard3* mRNA localization and MTOC correlative influence has not previously been demonstrated in fibroblasts as we do so here. Indeed our analytical approach was instrumental in revealing the presence of such enriched peripheral organization in polarized cells.

### Resolving spatial and temporal distribution of RNA and protein in a quantitative manner

We have shown our approach to be particularly suitable for the *de novo* identification of patterns of RNA spatial and temporal distribution and RNA-protein interdependent localization in a system, which has greater variability in terms of spatial localization and dynamics than developmental systems (Junker et al. 2014, Lecuyer et al., 2007). In particular, our analysis has demonstrated that micropatterning can be a valuable tool in studying RNA distribution as it allows us to reduce heterogeneity and isolate important modes of operation. The analytic tools we provide to identify sites of local interdependency between RNAs and proteins based on their dynamic patterns are readily applicable to other kinds of system, without requiring extensive manual annotation of visual features (Lecuyer et al., 2007). While previous approaches have identified broad classes of subcellular patterning in RNAs *de novo* via hierarchical clustering (Battich et al., 2013), lack of alignment limited the possible types of pattern identified and lack of integration with dynamic protein data limited the potential for drawing mechanistic and functional hypotheses from the patterns observed. Similar observations can be made in comparison to methods that have attempted to identify RNA patterns at the intercellular level, for instance in the zebrafish embryo (Junker et al. 2014), where hierarchical clustering based on spatial patterning alone is able to suggest similarity of function, but where integration of dynamic and other omics data is not attempted.

This has deep implications for projects such as the Human Cell Atlas (HCA). As noted above, several aspects of our approach are suitable as a basis for diverse kinds of spatially resolved omics (Crosetto et al., 2015), a key aim of the HCA. The quantitative nature of the analytic techniques introduced, autonomous image acquisition and automated features of our data processing make such techniques highly scalable to high-throughput studies and demonstrably in mammalian tissues. Particularly, the sensitivity of the approach to changes in localization under perturbation make the techniques suitable for inferring spatially organized regulatory networks (Crosetto et al., 2015) and can be combined with multiplexing techniques (Chen et al., 2015) to reveal dynamic changes in cell state. As well as interdependent mRNA-protein localization, our generalized clustering algorithms can be used to detect interdependent clustering dynamics between RBPs and different kinds of RNAs (lncRNAs, mRNAs, miRNAs), or interdependent protein-protein clustering patterns. Equally, our approach can be used to test various causal hypotheses for interdependent dynamics, such as characterizing RBP localization patterns, which act as determinants of lncRNA or mRNA localization (Lee et al., 2013), or building networks containing both RNA to protein and protein to RNA localization determinants. We believe that techniques such as those presented here will help to make possible the future development of such integrated approaches and contribute immensely to the HCA.

## EXPERIMENTAL PROCEDURES

### Cell culture and treatments

NIH/3T3 cells were grown at 37 °C to 100% confluence in DMEM/F12 medium supplemented with 10% FBS in a humidified atmosphere containing 5% CO_2_. Prior to micropatterning cells were serum-starved for 16 hr in DMEM/F12. For disruption of microtubule polymerization, nocodazole (Sigma) was added to a final concentration of 50 ng/ml to the medium, post removal of unattached cells and incubated for 3 or 5 hours before fixation of cells. Cytochalasin D (Sigma) was added to a final concentration of 1 mg/ml to the medium post removal of unattached cells and incubated for 1 h before fixation of cells.

### Cell micropatterning

Micropattern production was performed as previously described (Azioune et al., 2009). Briefly, glass coverslips were exposed to deep UV light using a UVO Cleaner (Jelight Company) for 5 mins. Cleaned coverslips were incubated with 0.1 mg/ml PLL-g-PEG (Surface Solutions) in 10 mM HEPES, pH 7.4 at RT for 1 hr. They were then rinsed once in PBS followed by one rinse in MilliQ water. The pegylated glass coverslips were then placed on a custom designed chromium photomask (Delta mask)(containing the desired micropatterns) and exposed to deep UV light for 5 mins. The patterned glass coverslips were then incubated with a fibronectin/fibrinogen-Alexa Fluor488 mixture (Life Technologies) in 100 mM NaHCO_3_, pH 8.5, at RT for 1 hr. The coverslips were then rinsed in PBS and used immediately for cell seeding. Serum-starved NIH/3T3 cells were seeded on the micropatterned surfaces at a density of 10,000 cells/cm^2^. After 30 mins, unattached cells were removed by gentle aspiration and replacement of the medium. Attached micropatterned cells were incubated at 37°C for 2 to 7 hours.

### RNA probes and reagents

Design and manufacture of RNA FISH probes for use in the single molecule FISH method were performed according to the protocol by (Raj et al., 2008). Multiple 20-mer oligonucleotide probes targeting the following mRNAs: *Gapdh*, *Arhgdia*, *β-Actin*, *Rab13*, *Pkp4* and *Pard3* were purchased (Biosearch Technologies). Each 20-mer contains a mdC(TEG-Amino) 3’ modification used to conjugate an NHS-ester ATTO-565 fluorescent dye (ATTO-TEC) to the probe. In brief, concentrated oligonucleotide probes were resuspended in 0.1 M Sodium tetraborate (Sigma) and mixed with resuspended 0.25 mg of the NHS-ester dye and incubated overnight at 37°C. This was followed by ethanol precipitation of the probes and purification by reverse phase HPLC on a XBRIDGETM OST C18 column to enrich for dye conjugated probes.

### Immuno-RNA FISH staining

For experiments utilizing the *Gapdh*, *Arhgdia*, *β-Actin* RNA probes: micropatterned NIH/3T3 cells were fixed in 3.7% formaldehyde for 10 min at 37°C followed by washes in PBS and overnight permeabilization in 70% ethanol at 4°C. For experiments utilizing the *Rab13*, *Pkp4*, *Pard3* RNA probes: micropatterned NIH/3T3 cells were fixed in pre-chilled methanol for 10 min, followed immediately by RNA FISH. The single molecule FISH method was modified from (Raj et al., 2008) to include immunofluorescence staining to detect the microtubule cytoskeleton. Cells were rehydrated in wash buffer (10% formaldehyde, 2X SSC) for 5 min. Hybridization was conducted overnight in a humidified chamber at 37°C in Hyb buffer (10% dextran sulfate, 1μg/μl E.coli tRNA, 2mM Vanadyl ribonucleoside complex, 0.02% RNAse-free BSA, 10% formamide, 2X SSC) combined with 50 ng of the desired RNA probe along with primary antibody - rat monoclonal anti-tubulin antibody (Abcam). Cells were then washed 2X (30 min at room temperature) with antibody wash buffer (10% formaldehyde, 2X SSC, anti-rat secondary antibody conjugated to Alexa Fluor 647 (Abcam)) followed by 1X wash with wash buffer. Cells were then incubated in equilibration buffer (0.4% glucose, 2X SSC) for 5 mins and counter stained with 1 μg/ml DAPI (4’,6-diamidino-2-phenylindole; Life Technologies). Coverslips were mounted in imaging buffer (3.7 μg/μl glucose oxidase and 1U catalase in equilibration buffer) and imaged.

### Immunofluorescence staining

Micropatterned cells were fixed in 3.7% formaldehyde for 10 min at 37°C, then washed with PBS followed by overnight incubation in 70% ethanol at 4°C. The cells were then washed with FBS followed by permeabilization for 10 min in 0.25% Triton-X at room temperature. Following this, the cells were washed thrice with PBS for 5 min each and incubated in blocking buffer (0.2 % BSA/PBS) for 30 min at room temperature. The cells were then incubated in the desired primary antibody solution (diluted in PBS) along with rat monoclonal anti-tubulin antibody (Abcam) to detect the microtubule cytoskeleton for 1 hr at room temperature. RhoGDI, Par3, β-Actin and Gapdh proteins were detected using rabbit polyclonal anti-Arhgdia (Santa Cruz), rabbit polyclonal anti-Pard3 (Abcam), rabbit polyclonal β-Actin (Santa Cruz) and rabbit polyclonal anti-Gapdh (Santa Cruz) respectively. Cells were then washed 3X with PBS following incubation with corresponding anti-rabbit secondary antibody conjugated to ATTO 550 (Rockland) together with anti-rat secondary antibody conjugated to Alexa Fluor 647 (Abcam) for 1 hr at room temperature. A further 3X wash with PBS was conducted followed by incubation in equilibration buffer (0.4% glucose, 2X SSC) for 5 mins and counter stained with 1 μg/ml DAPI (4’,6-diamidino-2-phenylindole; Life Technologies). Coverslips were mounted in imaging buffer (3.7 μg/μl glucose oxidase and 1U catalase in equilibration buffer) and imaged.

### Image acquisition

Most samples were imaged on a custom built spinning disk confocal Revolution XD system (Andor) comprising of a Zeiss Axio Observer.Z1 microscope with a 63X Plan-Apochromat objective (numerical aperture 1.4) and a cooled EMCCD camera (Andor iXon 897). Z-dimension positioning and control was accomplished by a piezoelectric motor (NanoScanZ, Prior Scientific). Images were captured using a custom developed algorithm based on ICY and μManager that allowed autonomous image acquisition (Figure S1A). In brief, the position of the micropatterns on the micropatterned surface were determined autonomously using the grid detection, alignment and calibration algorithm. This was then followed by sequential autonomous stepping through the micropatterned grid to determine the presence of a cell on the micropattern. If a single cell was detected on the micropattern surface by the algorithm then a *z*-dimension series of images was captured every 0.3 μm in four different fluorescence channels using emission filters for DAPI (DNA), Alexa Fluor 488 (micropatterns), ATTO 565 (mRNA/protein) and Alexa Fluor 647 (tubulin) and exposure times of 10 ms, 350 ms, 1 s (mRNA) or 500 ms (protein) and 350 ms respectively. A few samples being imaged on a custom built Nikon Ti Eclipse widefield TIRF microscope using a 100X N.A. 1.49 Nikon Apochromat TIRF oil immersion objective and equivalent fluorescent channels as above. After imaging, the data was processed using an automated background noise subtraction algorithm using ImageJ (Abramoff et al., 2004).

### Materials

Images were acquired for *Arhgdia*, *Gapdh*, *β-Actin*, *Pard3*, *Pkp4*, and *Rab13* genes. Tables 1, 2 and 3 below recapitulates acquisition conditions, techniques and number of acquired images in each series of FISH and IF data.

**Table 1.**
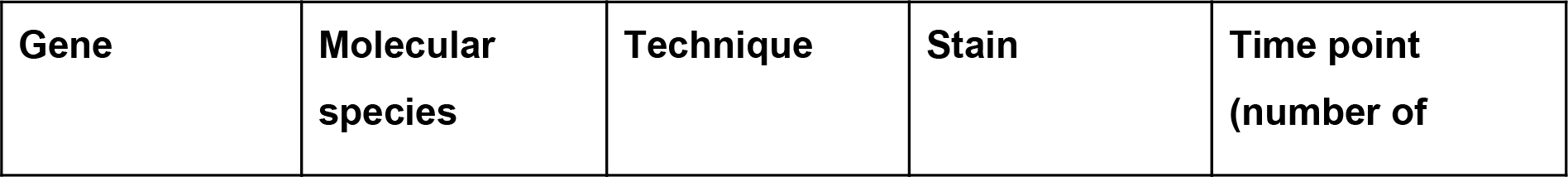
Image acquisition series characteristics and numbers for mouse fibroblast cells grown in micropatterned and standard cultures (indicated by *).

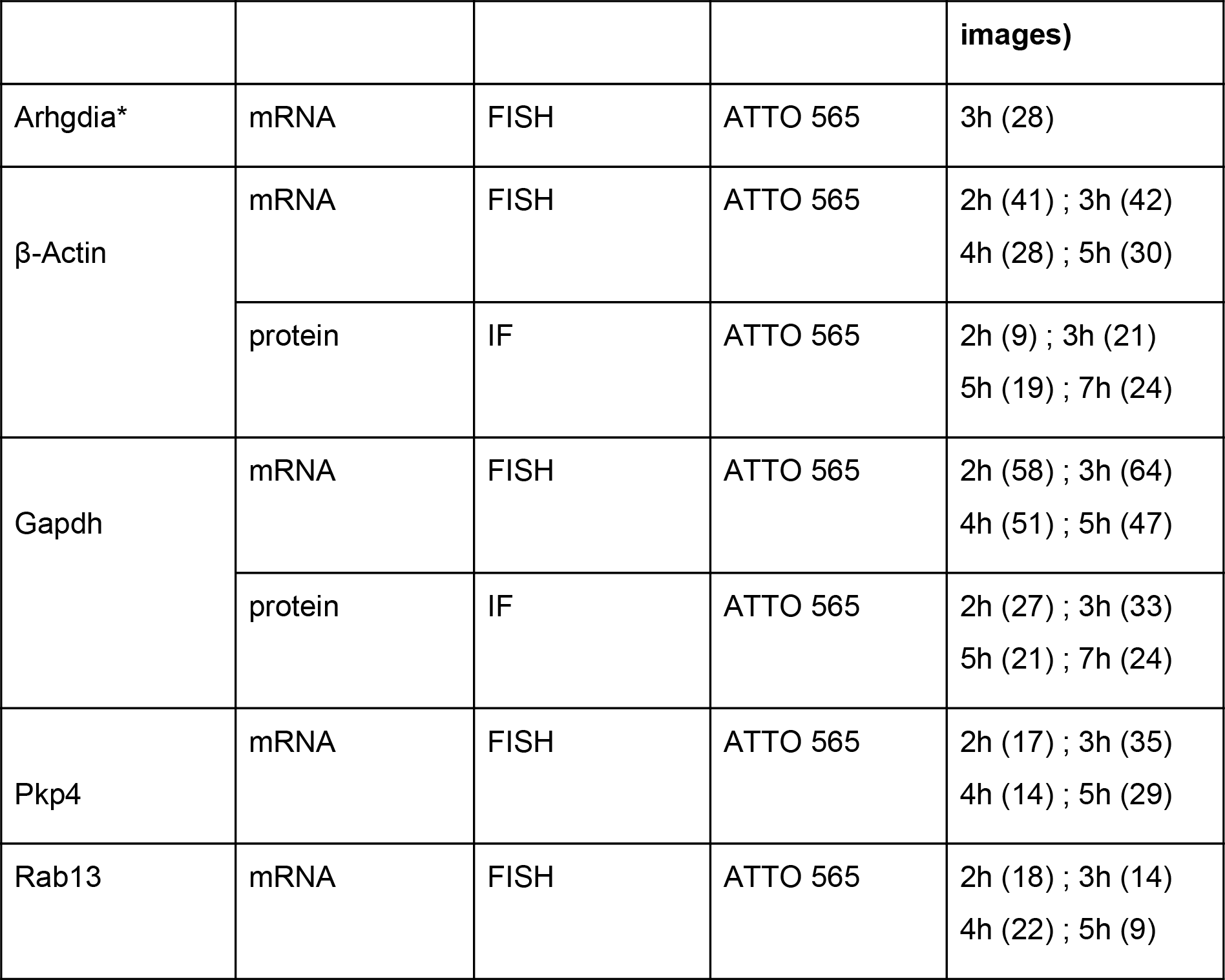

**Table 2.**
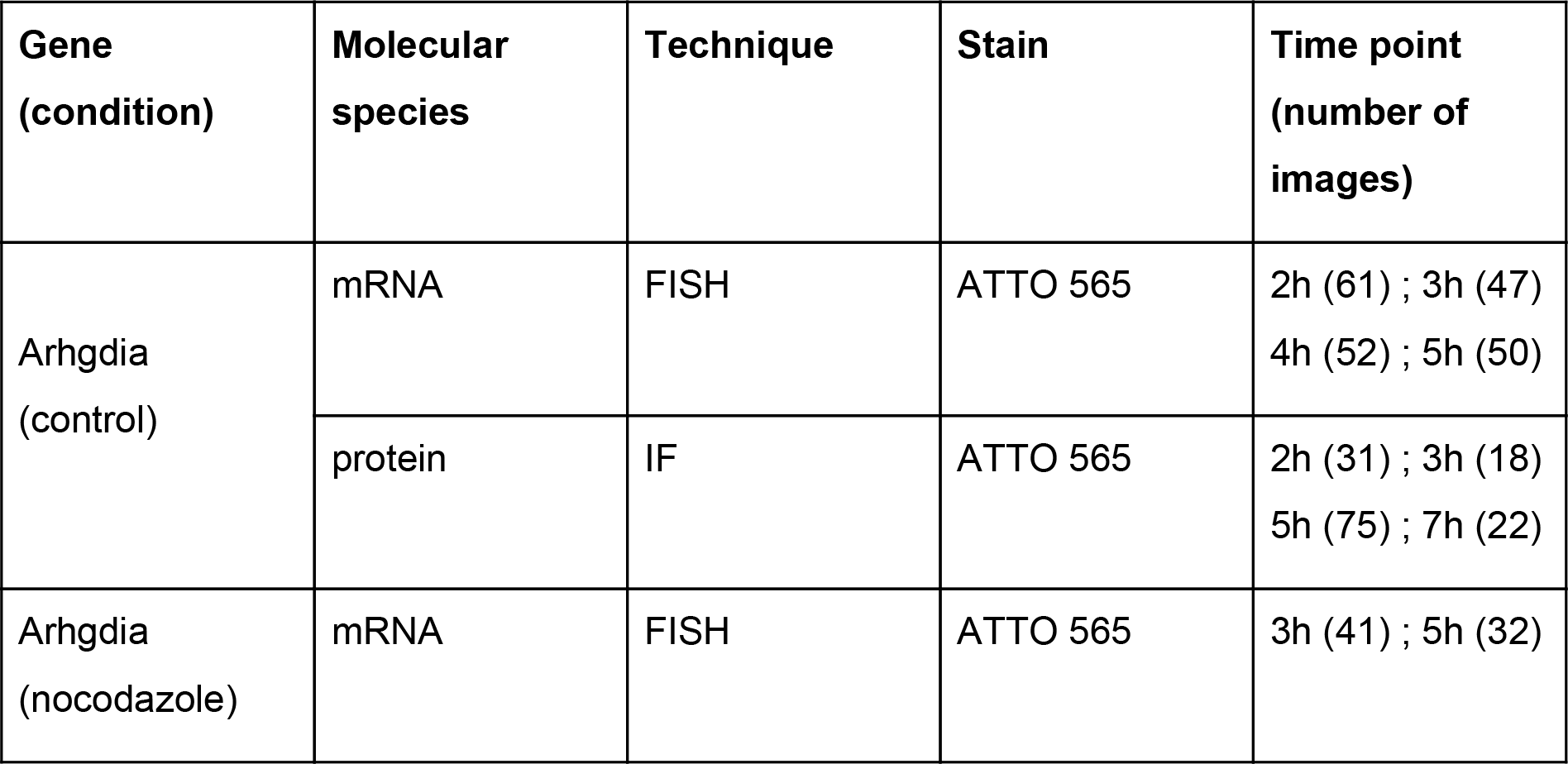
Image acquisition series characteristics and numbers for mouse fibroblast cells grown in micropatterned cultures in control and drug-disrupted conditions.

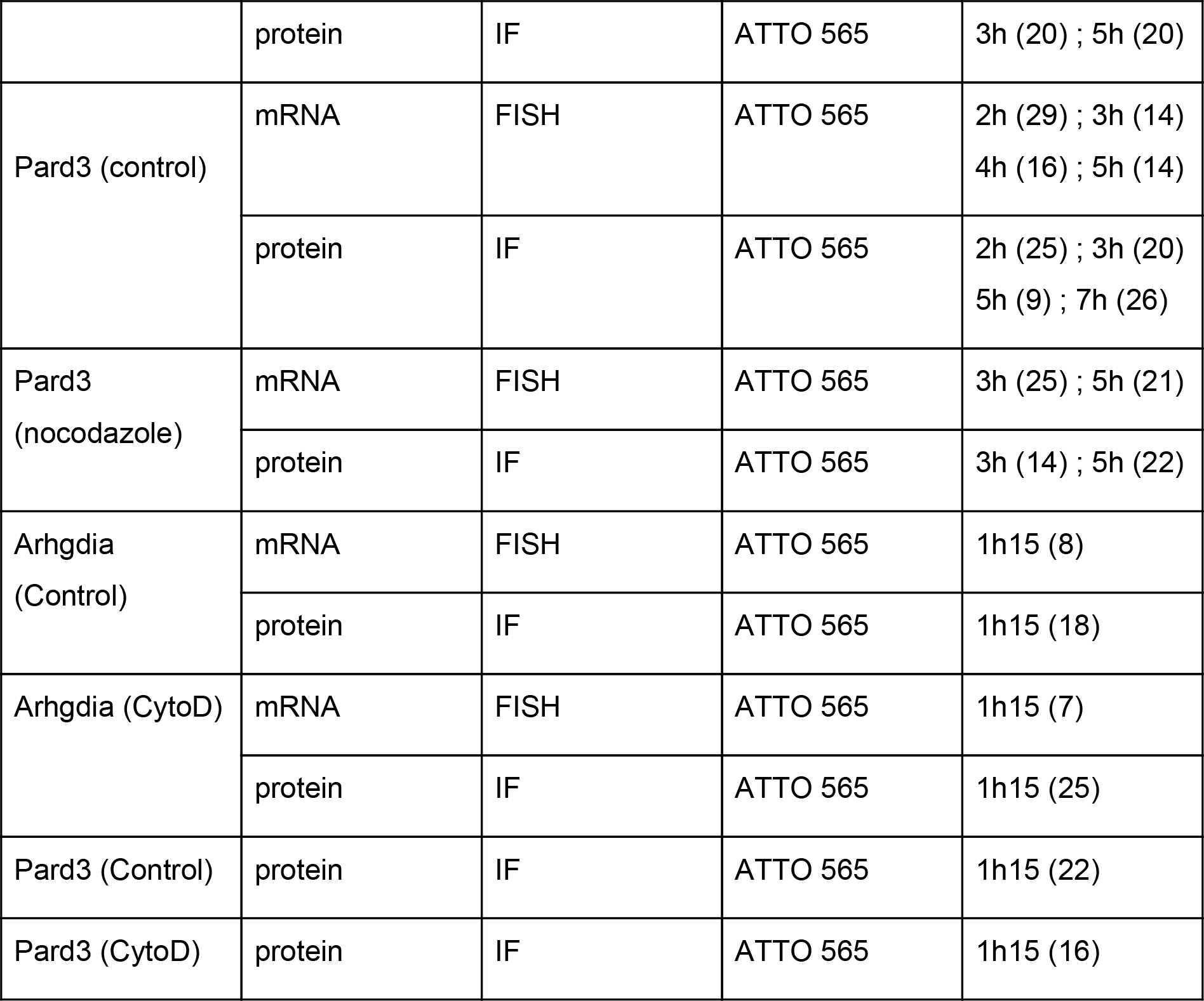

**Table 3.**
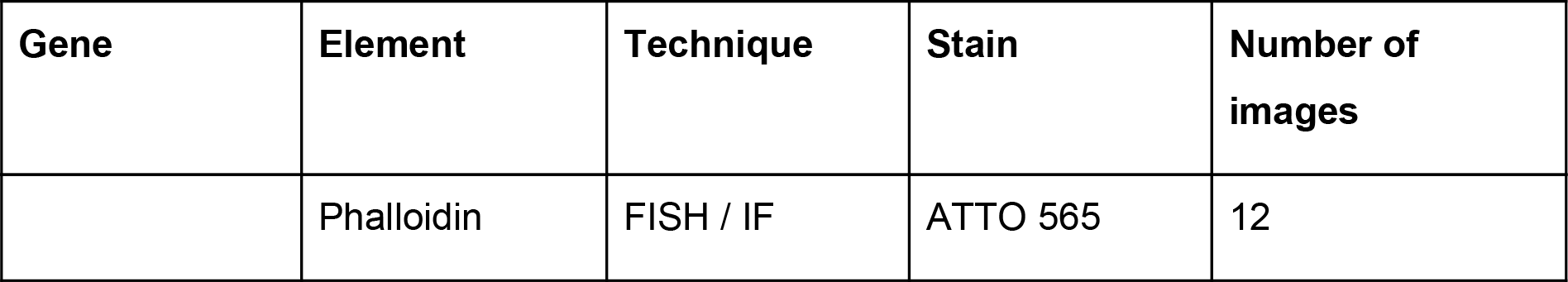
Image acquisition series characteristics and numbers for muscle cells.

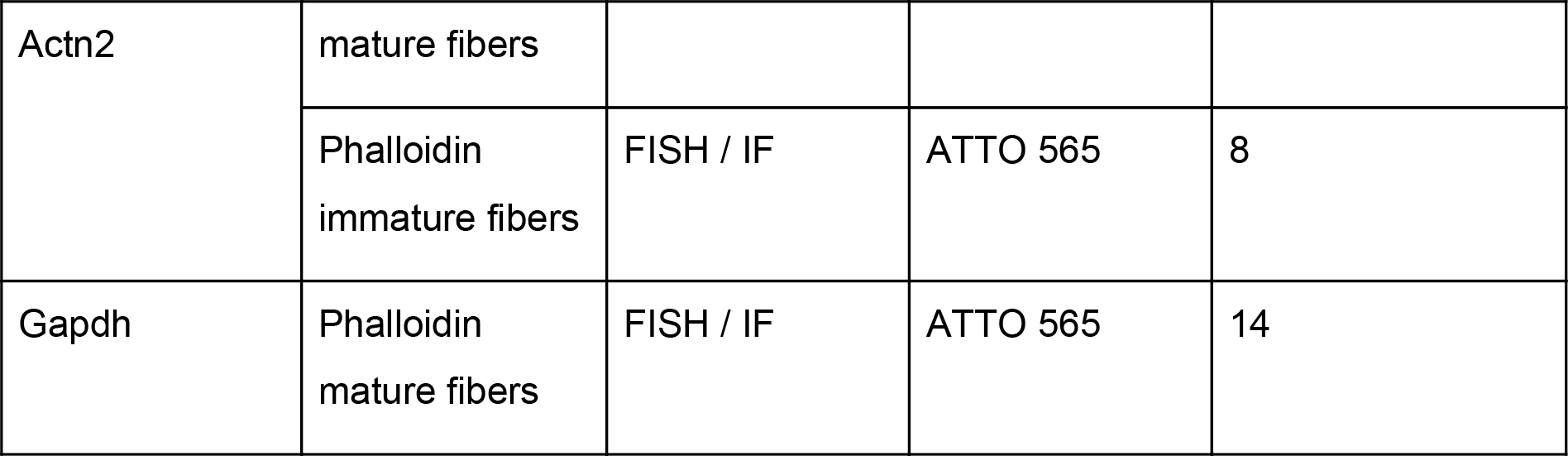

Myofibers were differentiated and fixed as previously described (pimentel et al + Roman et al NCB 2017).

Images were acquired for Actn2 and Gapdh genes. Acquisition conditions, techniques and number of acquired images in each series are recapitulated in Table 3.

For the purpose of image analysis we stained different cellular elements acquired at the same time as the FISH and IF signal, with differents staining as detailed in Table 4.

**Table 4.**
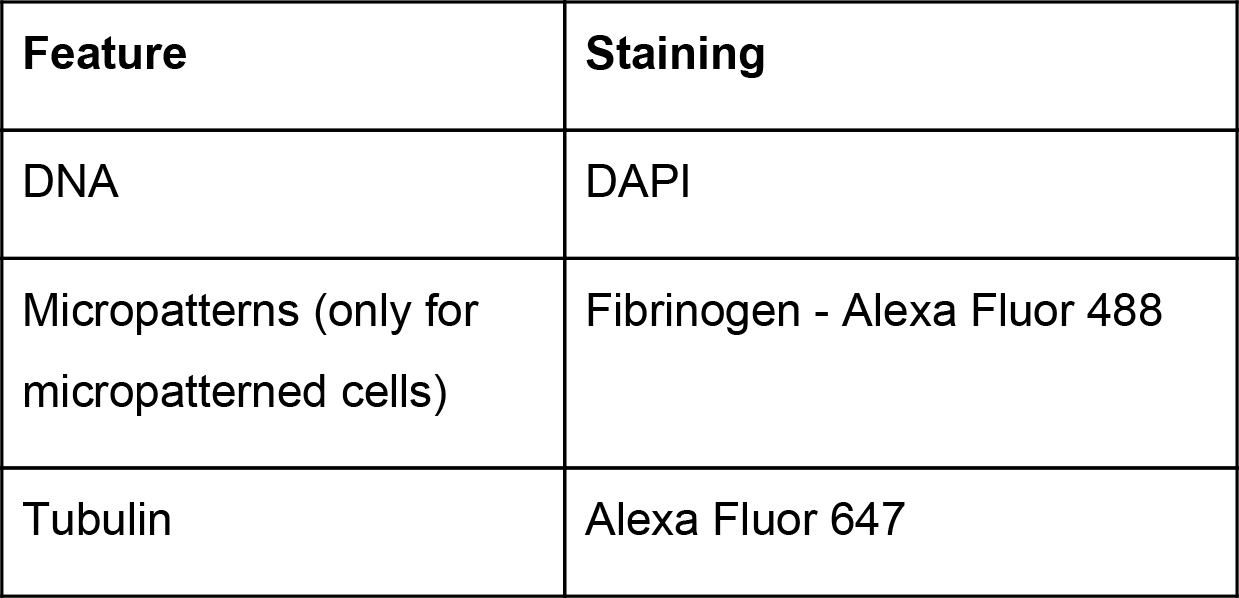
Summary of cellular features features stained in parallel to the FISH and IF signals.

| Feature | Staining |
| --- | --- |
| DNA | DAPI |
| Micropatterns (only for micropatterned cells) | Fibrinogen - Alexa Fluor 488 |
| Tubulin | Alexa Fluor 647 |

Images were acquired in the TIFF format. Our image processing pipeline transformed images into an HDF5 file (downloadable from the website www.dypfish.org).

### Image processing and statistical analysis

All computational analysis performed in the DypFISH project and described below were implemented in Python. Parameters for each of the algorithms that were used for each image acquisition series are available on the accompanying website (www.dypfish.org).

#### I. Primary image descriptors

Given the TIFF files, we first computed primary image descriptors for each image and stored them in an HDF5 file for each acquisition series.

##### (1). MTOC and nucleus centroid

FISH and IF images were manually annotated using the γ-Tubulin signal in order to obtain the coordinates (*x*, *y*) of the microtubule organizing center (MTOC). For the microtubule images the MTOCs were further annotated as being in the direction of the leading edge of the cell or not (Figure 1 Panel B). The nucleus centroid was computed as the geometric center of the nucleus mask (see below).

##### (2). Cell, nucleus and cytoplasm masks

Cell and nucleus masks were computed for all images (FISH and IF) using γ-Tubulin and DAPI signals, respectively.

For each image we obtained the maximum projection of the γ-Tubulin stained z-stack. A vignetting correction (Piccinini, 2013) is further applied to each resulting image individually by simply performing a pixel wise multiplication between each pixel value and the vignetting function. The detected cells being in the microscope’s focus, we assumed the optical center to be the center of the image and the intensity fall-off to be radially symmetric and the vignetting function is defined for each pixel *x*, *y* as *e*^−*d* / [(*w*/2)^2^*(*h*/2)^2^]^, where *d* = (*x* − *w*/3)^2^ + (*y* − *h*/2)^2^ and *w* and *h* are the image’s width and height, respectively. In a second time we perform contrast enhancement. Specifically, we apply histogram stretching by applying a linear normalization in order to stretch the interval of the intensities of a given image by fitting it to an another the [0, 255] interval.

First, we describe the procedure used for the detection of cell contours from the γ-Tubulin channel. We started by applying a local entropy filter to each pixel *i* as follows: 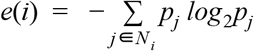, where *p_j_* is the proportion of pixels in the neighborhood *N_i_* having the same intensity as pixel *j*. The neighborhood size was chosen to be 30 × 30. For certain noisy image series we further applied a percentile thresholding tailored to each series of images on the resulting entropy histogram. As a last step, for all images we performed Canny edge detection, which detected edges by applying Sobel operators to the smoothed image, followed by hysteresis. However, the resulting edges were usually non-contiguous due to a weak γ-Tubulin signal or a high rate of noise. Consequently, we successively applied some mathematical morphological operators, such as dilation and closing, followed by erosion conventionally used to fill small gaps.

As the result of these steps we obtained a contiguous contour; small artificial white spots (artefact of the Canny filter) were eliminated by the previously used morphological operators. To this resulting image we applied the marching squares algorithm in order to obtain a 2D cellular segmentation mask, *M^cell^*(*x*, *y*), which is 1 for the cellular region and 0 otherwise.

For detecting nucleus masks *M^nucleus^*(*x*, *y*), the procedure was very similar using the DAPI signal, except that the local entropy filter was in most instances replaced by an Otsu filter, depending on the quality of the DAPI signal. Mathematical morphology algorithms were applied to neighborhoods ranging from 16 × 16 to 20 × 20 depending on the image acquisition characteristics (see for details on the www.dypfish.org).

Binary cellular and nucleus masks above were used to define a binary cytoplasm mask of the cell, *M^cytoplasm^*(*x*, *y*) = *M^cell^*(*x*, *y*)∧¬*M^nucleus^*(*x*, *y*)

##### (3). Zero level

An acquired image stack might contain irrelevant slices because the focal field of the microscope is outside the cell (above or below). To determine which slice contains the bottom of the cell and should be considered as the first relevant slice of the stack, we defined the *zero level* descriptor corresponding to the index of the slice having the maximum summed γ-Tubulin intensity. This zero level reference *z*-slice was used in further analysis such as e.g. the height-map computation or the degree of clustering.

##### (4). Height-map and cell volume

The height-map was built by segmenting each *z*-slice of a stacked image, which generated the 3D segmentation of the cell. It was performed for all FISH and IF images using the γ-Tubulin signal. Given a *z*-slice above the zero level we applied the cell mask detection procedure previously described, which defined a *z*-slice mask *M_z_*(*x*, *y*) with values corresponding to the height of the slice (*z*) within the mask and 0 outside. This set of slice masks defined the 3D representation, called height-map and denoted *h*(*x*, *y*) where the value at each coordinate (*x*, *y*) is the maximum over all slice masks, *max_z_*(*M_z_*(*x*, *y*)).

Based on the height-map we defined the cell volume 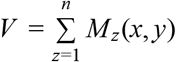 as the sum of volumes of all pixels within the height-map, where for each pixel *p* ∈ *M_z_*(*x*, *y*), its volume is *v*(*p*) = (1 ÷ 9.75μ*m*)^²^ × 0.3μ*m*, where 9.75 μ*m* is a size coefficient between pixel in μ*m*, and 0.3 μ*m* is the height of the slice ^1^.

##### (5). Protein intensities

Protein signal was computed for each immunofluorescence (IF) image as the sum of intensities across all z-slices and denoted as *I*(*x*, *y*).

##### (6). mRNA spot detection

To detect transcript positions from FISH data we used the ICY spot detector (Olivo-Marin et al., 2002). The detection was scripted so that for the images having *max*(*z*) ≤ 12 we used the following parameters: 2D wavelets and sensitivity 70 at pixel-scale 2; otherwise the parameters were set to: 1 pixel and 2 pixel length-scales with sensitivity 80. For cultured cells, as well as for CytoD micropatterned cell series, we have applied a custom-developed spot detection script (these images present a very high noise content preventing efficient use of ICY). First, we apply a background noise subtraction by using Sobel and Gaussian filters, successively. Second, we apply the white top-hat filter in order to enhance bright objects of interest (potential mRNA spots) on a dark background. Finally, we use the Laplacian of Gaussians filter for mRNA spot detection.

#### II. Secondary image descriptors

Based on the primary image descriptors we computed secondary descriptors that corresponded to per image statistics.

##### (1). Cytoplasmic total counts

Let us denote *M* the set of all mRNA spots for a given FISH image, |*M*| = *N*. The cytoplasmic total mRNA descriptor was calculated as the number of transcripts within *M^cytoplasm^*, that is *T_mRNA_* = | {*m* ∈ *M* | *M^cytoplasm^*(*x*, *y*) = 1 |. The cytoplasmic total IF intensity is the summed IF intensity across the *M^cytoplasm^* region for protein images:

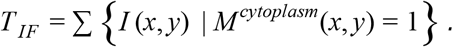

##### (2). Peripheral distance map

For a given image, the peripheral distance map corresponds to a collection of peripheral masks based on *M^cytoplasm^*(*x*, *y*), where the width of the periphery varies as a proportion of the cytoplasmic radial distance. We segmented *M^cytoplasm^*(*x*, *y*) into 100 isolines from the nucleus contour to the periphery by projecting a ray from the nucleus centroid to the cell border, which was then segmented in 100 equidistant points. The 100 isolines were then built by constructing polygons that connect 360 points (one ray per degree). These isolines define a symbolic distance map *D*, where *D*(*x*, *y*) is the isobar value for (*x*, *y*) corresponding to the “distance” from the nucleus, with 100 at the nucleus and 0 at the cell edge. Given a fixed percent *p* between 0 and 100, the mask *M^periphery^* (*x*, *y*, *p*) is 1 for *D*(*x*, *y*) < *p* and 0 otherwise. Hence, the periphery mask for a given *p* contains a strip at the cell edge whose width is a fixed proportion of the radial distance.

#### III. Statistical analysis

Primary and secondary image descriptors were used to compute statistics for image acquisition series and to compare them.

##### (1). Peripheral fraction and enrichment

Based on the cell masks, we calculated the peripheral fraction of mRNA and proteins at a given percent *p* of the radial distance. This fraction is defined as the ratio of the transcript counts (respectively, summed IF intensities) across the *M^periphery^* (*x*, *y*, *p*) and *M^cell^* regions.

Each mRNA spot located at (*x*, *y*) has its corresponding peripheral distance, defined by the value of the distance map *D*(*x*, *y*) at the same coordinate, which provides a mapping *d* : *M* → *D*. This defines a vector *C* = (*n*_1_, … *n*_100_) containing the counts of mRNAs at distances *i* ∈ [0, …, 100] from the cell edge normalized by the total number of mRNAs, that is *n_i_* = |{*m* ∈ *M* | *d*(*m*) = *i*}| / *N*. The mRNA fractions for each gene and for each isobar were defined as vector 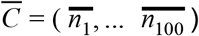 of means over all FISH images for this gene over all time points. The peripheral fraction of mRNA for a given gene and given *p* was then computed as 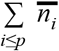.

##### (2). Volume corrected noise measure

In order to measure gene expression noise while accounting for cell volume, we computed the volume corrected noise measure *Nm* for micropatterned and standardly cultured cells. It was calculated following the approach of a previous study (Padovan-Merhar et. al. 2015):

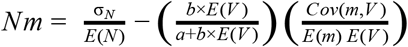

where *N* is the total mRNA count, *V* is the cell volume, *a*, *b* are the offset and slope of the least-squares best-fit linear regression of *E*(*N*) on *V*, and σ, *E* and *Cov* are the notations for standard deviation, expectation and covariance, respectively.

##### (3). Cytoplasmic spread

Cytoplasmic spread is a statistics that measures how evenly a molecule is spread across the cell. For the mRNAs it corresponds to the average distance from the nucleus centroid of cytoplasmic mRNAs normalized by the total cytoplasmic cell spread. For the protein intensities it is the expected distance from the nucleus centroid according to the protein cytoplasmic intensity distribution, normalized by the cell spread. In both cases the statistics takes value 1 when the molecules are evenly distributed across the cytoplasm.

First we defined the cytoplasmic cellular spread *S* as the average distance of a cytoplasmic voxel from the nucleus centroid in the *x* − *y* plane:

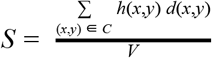

where *C* is the set of all coordinates (*x*, *y*) for which *M^cytoplasm^*(*x*, *y*) is 1, *h* is the previously defined height map, *d*(*x*, *y*) the 2D euclidean distance of pixel (*x*, *y*) from the nucleus centroid, and *V* is the cell volume.

The mRNA cytoplasmic spread is then defined as *M* = *m* / *S*, where m is the mean 3D distance of all cytoplasmic transcripts, that is those where *M^cytoplasm^*(*x*, *y*)= 1, from the nucleus centroid.

The protein cytoplasmic spread is defined as:

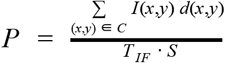

where *I*(*x*, *y*) is the summed IF signal intensity at a given coordinate (*x*, *y*) and *T_IF_* is the cytoplasmic total count of the IF signal intensities.

##### (4). Cell quantization

In order to compute localisation statistics over multiple micropatterned images compatibility of these images is required. We have chosen the MTOC position to be the reference point for the 2D cell geometry.

###### (A) 2D quantization: quadrants and per quadrant statistics

As shown on the schematic in Figure 3, panels A and B, given a cell mask *M^cell^*(*x*, *y*), we generate the tessellation of the image by centering two orthogonal axes at the nucleus centroid and rotating them over 360 degrees, each position of these axes defining a partition of the cell mask into four quadrants, one of them containing the MTOC, *Q_M_*. We retained the orientation that yields the maximum mRNA count within a quadrant containing the MTOC, that is *max_d_*(*T_mRNA_* = |{*m* ∈ *M*|*M^cytoplasm^*(*x*, *y*) = 1 ⋀ *Q_M_*(*x*, *y*) = 1}), *d* ∈ [0, 359]. The resulting fours quadrants *Q*_1_, *Q*_2_, *Q*_3_, *Q*_4_ are numbered so that *Q*_1_ always corresponds to *Q_M_* and the the remaining three quadrants are numbered in the clockwise fashion.

For protein intensities, quadrants are defined in the similar fashion using *T_IF_*. Definition of cell mask partitioning in quadrants *q*, *q*_2_, *q*_3_, *q*_4_ enables cell’s quantization in 2D in terms of per quadrant statistics of mRNA and protein signal. Quadrants’ respective areas are denoted by *a*_1_, *a*_2_, *a*_3_, *a*_4_. We denoted by *t_i_* the total number of mRNA spots falling in *q_i_* in the case of FISH data, or the summed intensity across *q_i_* in the case of IF data.

Then the local mRNA density was computed as the relative concentration *c_i_* of mRNA in quadrant *i* and is defined to be 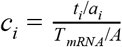, where *A* is the cell mask area. In the case of protein signal we replaced *T_mRNA_* by *T_IF_*.

###### (B) Fine-grained quantization

In the same fashion as for the peripheral distance map we defined an additional subdivision of the cellular mask in isolines, their number being defined by the percent *p*. Given the previously defined quadrants, we further subdivided each of them in 2, yielding the tessellation in 8 parts that divide the circle in 45 degree sectors. Using the isolines and the 8 sectors we quantized the cell masks into 8 × *p* segments organized in a concentric fashion starting from the nucleus towards the cell periphery (see the schematic in Figure 4, panel A). Quantization for thus obtained segments was computed in the same fashion as for quadrants, resulting in a 8 × *p* vector of per segment signal concentration statistics for each cell, that we denoted *C* = (*c_i_*).

###### (C) 3D quantization

Cell mask’s tessellation into quadrants as defined in III.4.A (axes position) is projected onto each *z*-slice, thus yielding the cell’s partition into four 3D quadrants *Q*_1_, *Q*_2_, *Q*_3_, *Q*_4_, their respective volumes being denoted by *v*_1_, *v*_2_, *v*_3_, *v*_4_. The volume of each quadrant is calculated as the sum of volumes of pixels within it using the same coefficients as for the cell volume.

We denoted by *t_i_* the total number of mRNA spots falling in *Q_i_* in the case of FISH data, or the summed intensity across *Q_i_* in the case of IF data.

Then the relative concentration *c_i_* of mRNA in quadrant *Q_i_* is defined to be 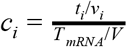. In the case of protein signal we replaced *T_mRNA_* by *T_IF_*.

##### (5). MTOC polarity index

We defined a polarity index *PI_M_* ∈ [− 1, 1], called the *MTOC polarity index*, that measures the enrichment of mRNA or protein signal for a given image acquisition series in the vicinity of the MTOC location.

For the set *S* of images from an acquisition series under study, we denoted by *S_M_* = {*S_i_*} and *S*_¬*M*_ = {*S_j_*} the sets of all MTOC containing quadrants and quadrants that do not contain the MTOC, respectively. Intuitively, the MTOC polarity index measures how frequently the concentration within the MTOC quadrants is higher than in the non-MTOC quadrants. Formally it is defined as follows:

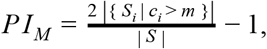

where *m* is the median of signal concentrations *c_j_* for all quadrants in *S*_¬*M*_.

Positive values of *PI_M_* imply MTOC correlated enrichment of RNA transcripts or proteins, negative values imply enrichment away from the MTOC and a value of zero implies no detectable correlation.

Statistical relevance of *PI_M_* is measured using the null hypothesis that ∀_*i*_, *c_i_* = *m*, which corresponds to the complete spatial randomness. Under this hypothesis the population value of *PI_M_* is 0. However, we have shown in (Warrell et al., 2016) that the empirical distribution of *PI_M_* follows the binomial distribution asymptotically. Thus, the binomial test was used to evaluate the statistical relevance of *PI_M_* for a given set of images.

##### (6). mRNA / protein distribution profile

In order to define a spatial distribution profile of mRNAs and proteins for images acquired at a given time point, we used the fine-grained quantization of the cells (see paragraph IV.4.B). A single vector was computed at each time point by averaging across the pool of acquired images, hence estimating its expected value at that time point. Recall, that for each cell we computed a vector *C* = (*c_i_*) of per segment signal (mRNA or protein) concentration statistics. Then for a given time point we computed a mean spatial profile *C̅* representative of this time point by averaging all *C_i_* for this acquisition series.

##### (7). Temporal interaction score

The goal of this analysis is to measure the interdependence between the mRNA and protein dynamics. To do this, we defined the *Temporal Interaction Score* (TIS) as a correlation between mRNA and protein spatial distributions for image acquisitions at several time points. TIS for a given mRNA-protein pair is calculated based on mRNA and protein distribution descriptor vectors 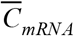 an 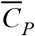.

TIS can be calculated for any measure of correlation between mRNA and protein distributions, which allowed us to examine the interdependence of molecule’s dynamics within specifically defined subcellular regions.

More formally, we supposed a 2-measure discrete time process Φ, containing Observations 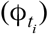 and 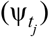 at time points *T*_1_ = {*t_i_*} and *T*_2_ = {*t_j_*}, respectively. Then, the (empirical) temporal interaction score, 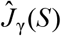 is defined for pairs of data 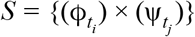 at time points *T*_1_ and *T*_2_ using a similarity function γ in the following way:

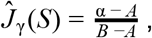

where α is the rank-sum 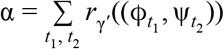, *t*_1_ < *t*_2_, *r*_γ_ is the rank of the tuple where the order is given by the γ function, γ function is the similarity between pairs of observations is computed as 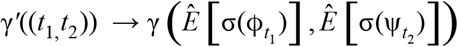. Constants *A* and *B* ensure that 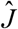 lies between 0 and 1 and are defined using the notion of ‘forward-leading’ time point pairs.

The ‘forward-leading’ set *S*_1_ is defined as *S*_1_ = {(*t*_1_, *t*_2_) ∈ *T*_1_ × *T*_2_ | *t*_1_ < *t*_2_}, and its complement *S*_2_ contains all pairs of time such that *t*_1_ ≥ *t*_2_. We define constants *A* and *B* as *A* = (|*S*_1_|/2)(|*S*_1_| + 1) and *B* = (|*S*_1_||*S*_2_|/2) (|*S*_1_||*S*_2_| + 1)(|*S*_2_|/2)(|*S*_2_| + 1). Thus, 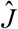 can be understood as the rank-sum of the similarities γ′(*t*_1_,*t*_2_) across all forward-leading’ time point pairs normalized by *A* and *B* to lie between 0 and 1.

Consequently, observations from Φ are ranked in the ascending order according to the value of the similarity of some statistics σ between ϕ and Ψ.

In practice, for the analysis of mRNA / protein interactions, we considered computing the TIS for a 2-measure discrete time processes Φ in which ϕ is a point process and Ψ a general random measure (representing mRNA locations and protein concentrations respectively), *T*_1_ = {2, 3, 4, 5} and *T*_2_ = {2, 3, 5, 7} (the discrete time points representing time in hours), σ(.) is the normalized histogram over a fixed finite set of voxels as described in the section III.6, and γ is the Pearson Correlation Coefficient between two histograms. We note that σ(.) can represent a histogram based on a particular quantization of cells and the histogram can cover the whole cell (forming a global TIS).

Moreover, we evaluated whether there is a deterministic influence of the mRNA distributions on the protein distributions of late time points against a null hypothesis of dynamic equilibrium. We used the empirical temporal interaction score 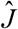 to identify non-stationary dynamics given pairs of mRNA and protein data *S*, and we tested whether the null hypothesis (that Φ is in a dynamic equilibrium) can be rejected.

In (Warrell et al., 2016) we have shown that 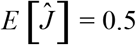 under the null hypothesis that the process Φ is at steady-state, and further that its distribution can be characterized up to a dependence on the ranking function. From the rank-sum of the similarities γ′((*t*_1_, *t*_2_)) across *S*_1_ defined previously, an exact permutation test can be derived to calculate significance levels for a given value of 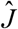 and a steady-state null hypothesis.

##### (8). Degree of clustering (Ripley-K)

The degree of clustering statistic has been previously introduced based on the framework of point processes by (Lee et al., 2013). It is a unitless measure that can be used to compare clustering between different molecules and conditions. In (Warrell et al., 2016) we generalized this definition to the framework of continuous random measures, which allows us to calculate the degree of clustering for both FISH and IF data, the former being modelled as point processes, and the latter modelled as a continuous-valued random measure. Our generalized algorithm for calculating the degree of clustering is summarized below. For theoretical considerations please see (Warrell et al., 2016).

A classical tool for the point process analysis is the Ripley’s K function defined as the mean number of events that occurred inside a ball of radius *r* around a randomly selected event normalized by λ, the number of events per unit area (Ripley, B. D. 1977). A classical estimator of the Ripley’s K function can be defined as in (Chui et al., 2013, Ripley 1977):

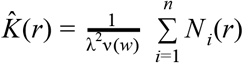

where *N_i_* is the number of event points (mRNA transcripts) in a ball of radius *r* centered on *i*, λ is the density, ν is the volume or area (in 3D and in 2D, respectively) of the observed region *w*, and *n* is the number of points.

We normalized 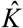 under a homogeneous Poisson process, which is commonly known as Ripley’s H function 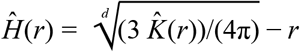, *d* ∈ {2, 3} where *d* is equal to 3 in the case of volume-based computation and 2 in the case of 2D.

This in turn makes it possible to define the clustering index *H*^*^ as an estimator of 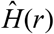 by comparing the Ripley’s H function calculated empirically to its distribution under complete spatial randomness (CSR):

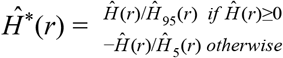

where 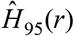 and 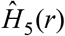 are the 95th and 5th percentiles respectively of 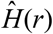.

CSR is modeled using random permutations of actual data points (100 times in our study), which enabled us to compute the 95% and 5% confidence bounds of CSR. Spatial clustering is considered to be significant at radius *r* if the computed 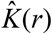 is over the upper (95%) or lower (5%) bounds of the random distribution.

In (Warrell et al., 2016) we have introduced a convolution-based 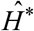 estimator based on the exact permutation-test. This estimator normalizes 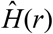 so that 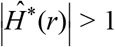 only when 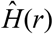 falls outside the 95% confidence interval for a homogeneous Poisson Process. Moreover, we have shown that our permutation test using the convolution-based estimator reduced to the clustering index estimator used by (Lee et. al. 2013) for the point process case. This enabled the implementation of a common consistent computational framework for both point and continuous processes. The degree of clustering 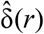 is then defined as the area of 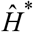 above 1, that is 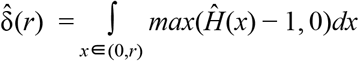.

Within this unified framework we can use the same computational approach for mRNA data as in (Lee et al., 2013) to compute the degree of clustering. Below we define the specific procedure for its computation in the case of protein data.

Cells are quantized into voxels *V*_1_,…, *V_n_* where each voxel t the value of the observed quantity ϕ(*V*_1_), …, ϕ(*_n_*) in the case of 3D analysis (or into pixels in the case of 2D). We denoted by *I* an array in which each element corresponded to the intensity value for each voxel.

The convolution-based estimator 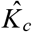 can be computed using the following formula:

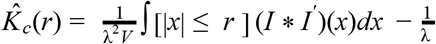

where λ is an estimate of average intensity per unit volume, [.] is the indicator function that is 1 for a true statement and 0 otherwise, *I*^′^(*x*) = *I*(− *x*), * is the convolution operator, and *V* is the volume of the window over which the cell is observed. In practice, given the fact that the cell thickness is quite low, we can approximate the 3D convolution by a 2D convolution.

Thus we have a common computational framework to evaluate the presence or absence of clustering for both mRNA and protein data.

#### IV. Additional methods for muscle data analysis

In this section we report adaptations of the methods presented in sections I, II and IV to the case of muscle cells (see Table 3).

##### (1). Cell and nucleus masks, nucleus centroid

Cell and nucleus masks were computed for all muscle images, using γ-Tubulin and DAPI signals, respectively, using the same general principles as in section II.2. As these acquisition series benefit from a better segmentation, only Otsu threshold method was necessary to obtain the binarized images.

After these steps, we obtained a contiguous contour with small white spots due to the noise in the images. We applied mathematical morphological operators such as dilatation and closing to get a full mask of the muscle cell *M^cell^*(*x*, *y*). The nucleus mask *M^nucleus^*(*x*, *y*) detection followed the exact same steps and parameters as for the micropatterned images - the nucleus centroid was computed as the geometric center of the nucleus mask.

##### (2). mRNA and protein signal detection

mRNA spots detection was done using ICY spot detector (Olivo-Marin 2002) to find transcript positions, the parameters were set to: 1 pixel and 2 pixel length-scales with a fixed sensitivity of 80.

##### (3). z-lines’ masks

The main component of z-lines is the Alpha actinin protein. To facilitate the analysis, we have defined an additional secondary descriptor computed from the Phalloidin signal, called z-lines mask *M*^*z*−*lines*^(*x*, *y*).

For each *z*-slice of each image we performed the contour detection for the z-lines. First, we applied a vertical Sobel operator, which detected the vertical edges of an image, followed by a Gaussian kernel to smooth artifacts of the Sobel filtering and reinforce the z-line signal. An Otsu binarization was then processed. As a result we obtained a set of z-lines masks 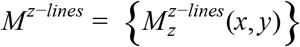 (Figure 6).

For further analysis we restricted the cell to *z*-slices containing more than 25 mRNA spots (to avoid false positives due to high noise). Notice that the spots falling in the eliminated slices were also excluded from the analysis.

We defined an additional descriptor called z-line spacing *Z* that represented the median spacing between 2 lines. For each *z*-slice at each *y* coordinate we computed all the distances *d*((*x_i_*, *y*), (*x_j_*, *y*)) where *x_i_* and *x_j_* were 2 consecutive z-lines contours. *Z* was defined as the median of all *d* for all acquired cells. For our data *Z* = 15 pixels.

##### (4). z-line mRNA distance profile

In order to evaluate the mRNA clustering in the vicinity of z-lines, we computed the average 2D euclidean distance of mRNA spots to their nearest z-line for mature and immature cells.

Using the *M*^*z*−*lines*^(*x*, *y*) masks, we computed for each mRNA spot *m* ∈ *M* positioned at (*x*, *y*, *z_m_*) the minimal 2D Euclidean distance to a z-line falling within a disk of radius *Z*. This computation was performed within the 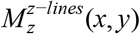 z-mask such that *z* = *z_m_*. If *m* fall within 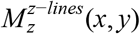, then the minimal distance was set to 0. Thus for each image we obtained a *D* = {*d*}, |*D*| = *N* the set of all minimal distances between mRNA and z-lines. In turn this allowed us to define for each image a count vector δ = (δ_*i*_) of size *Z* where δ_*i*_ is the number of mRNA spots at each distance *d* = *i*, *d* ≤ *Z* normalized by *N*.

For a given image acquisition we defined its z-line mRNA distance profile to be 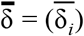, where 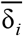 is the median of all δ_*i*_ (Figure 6, panel B).

##### (5). Cell quantization

Given that muscle cells contained more than one nucleus, each cell mask was restricted to be between two consecutive nuclei centroids as shown on the schematic in Figure 6 (panel C). Given a cell mask *M^cell^*(*x*, *y*), definition of cell mask tessellation in *n* vertical segments *q_i_*… *q_n_* enables cells quantization in 2D in terms of per segment statistics of mRNA concentrations. Quantization was performed with *n* = 20 and *n* = 80, see results in Figure 6 (panel C).

##### (6). mRNA spatial distribution

To estimate mRNA clustering along muscle cells, we computed local mRNA density for each cell using the cell quantization introduced in section V.5. We denote by *t_i_* the total number of mRNA spots falling in a given *q_i_*. Then the local mRNA density is computed in the same way as for fibroblast cells (see section III.1) as the relative concentration *c_i_* of mRNA in *q_i_* and is defined to be 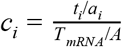, where *A* is the cell mask area. We produced distribution plot and heatmap representing mRNA local density between two nuclei (Figure 6 panel C).

## AUTHOR CONTRIBUTIONS

M.M.M. designed the overall study with contributions from A.F.S, R.B., M.N. and J.H.W.. A.F.S. designed and carried out experiments, collected and analyzed data and co wrote the paper. R.B. designed and carried out experiments, collected and analyzed data and co-wrote the paper. M.N., E.B and B.D coded and implemented the analytical algorithms and co-wrote the paper. J.H.W. and M.M.M. developed the beta versions of DypFISH. S.D. and J-C.O-M. developed and implemented autonomous image acquisition and data collection for microscopes. J.S. helped with the design of the study. R.B, A.F.S., J.H.W., J.S and M.M.M. discussed and edited the paper. M.M.M. designed experiments, analyzed data, supervised the study and co-wrote the paper.

## ACKNOWLEDGMENTS

We thank all members of the Gene Expression and Biophysics Laboratory (Mhlanga Lab). We thank N. Crosetto, and members of the L. Pelkmans lab, members of the J-C. Olivo-Marin lab, K. Schauer and members of the B. Goud lab for comments on this manuscript. A.F.S is a recipient of the postdoctoral fellowship from the Claude Leon Foundation, South Africa. This research has been supported by the following grants all to M.M.M. V2YGE81 (to A.F.S.) and PG-V2KYPO7, TA 2011 011 from the Council for Industrial and Scientific Research (CSIR, South Africa) and by a grant from the Emerging Research Area Program of The Department of Science and Technology (DST, South Africa) Department of Science & Technology Centre of Competence Grant, SA Medical Research Council SHIP grant, CSIR Parliamentary Grant and M.M.M. is a Chan Zuckerberg Investigator of the Chan Zuckerberg Initiative - Human Cell Atlas program.

## SUPPLEMENTARY FIGURE LEGENDS

**Figure S1, related to Figure 1.**
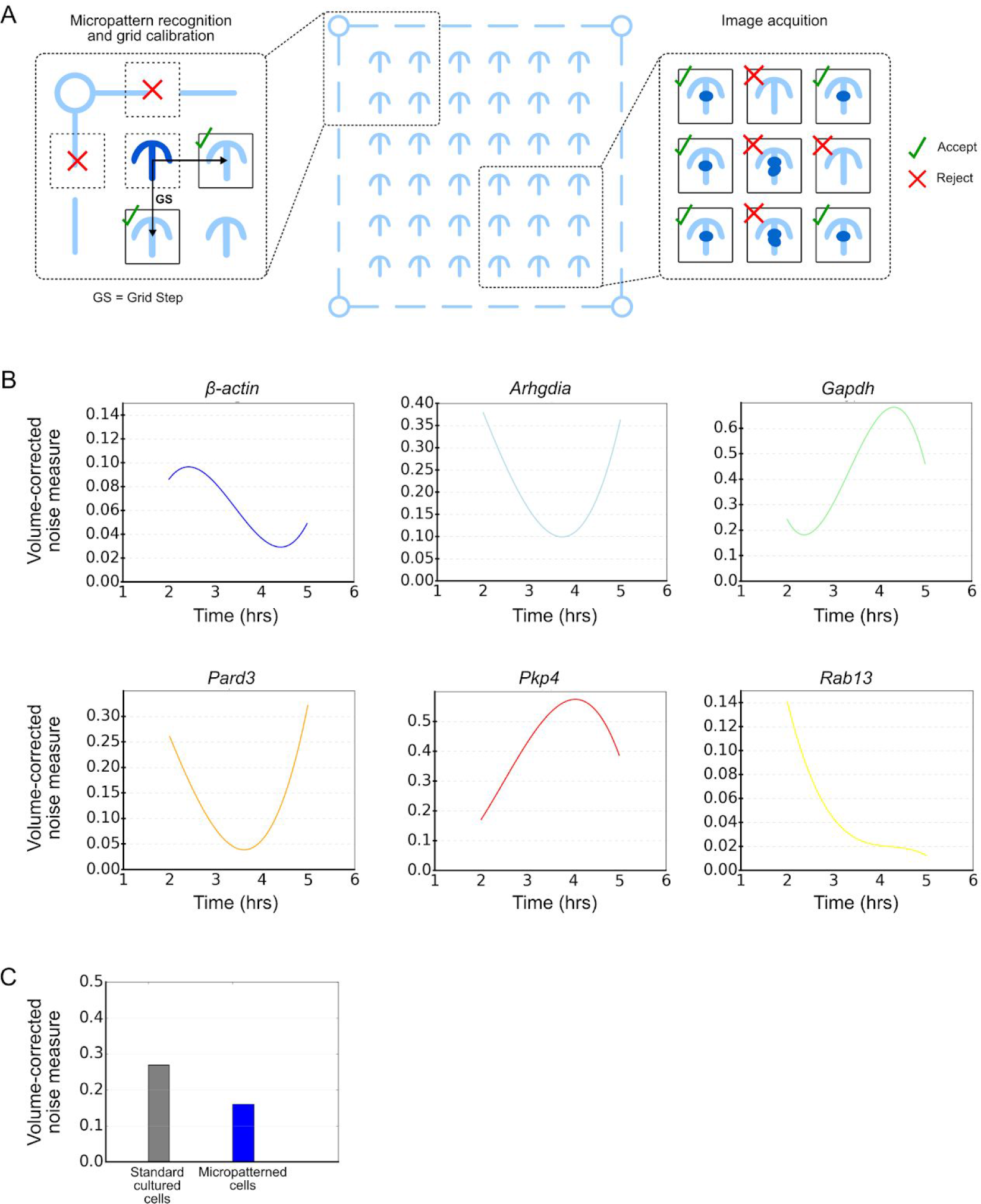
Automated image acquisition and effects of micropatterning on noise. (A) Each coverslip was micro-fabricated to contain multiple 12 by 12 grids of micropatterns to which the cells adhered facilitating the development of an algorithm to automate the process of image acquisition. The algorithm initially performed grid calibration in which the location of the upper-left micropattern was automatically detected, followed by grid orientation and grid step size determination (left). Images were then collected at each grid position across the full 12 by 12 grid. A 2-class support vector machine was trained to classify cells which have grown normally on the micropatterns versus micropatterns containing no cells, multiple cells, or cells which have failed to fill the micropattern, allowing the automatic rejection of grid positions which cannot be used (right). (B) Comparing the volume-corrected noise measure (Padovan-Merhar et al., 2015) across time for 6 mRNAs. Cubic splines are fitted to the values measured at 2, 3, 4 and 5h time-points. © The stochasticity which remains after correcting for the linear relationship between cell-size and transcript number using the volume-corrected noise measure (Padovan-Merhar et. al. 2015).

**Figure S2, related to Figure 3.**
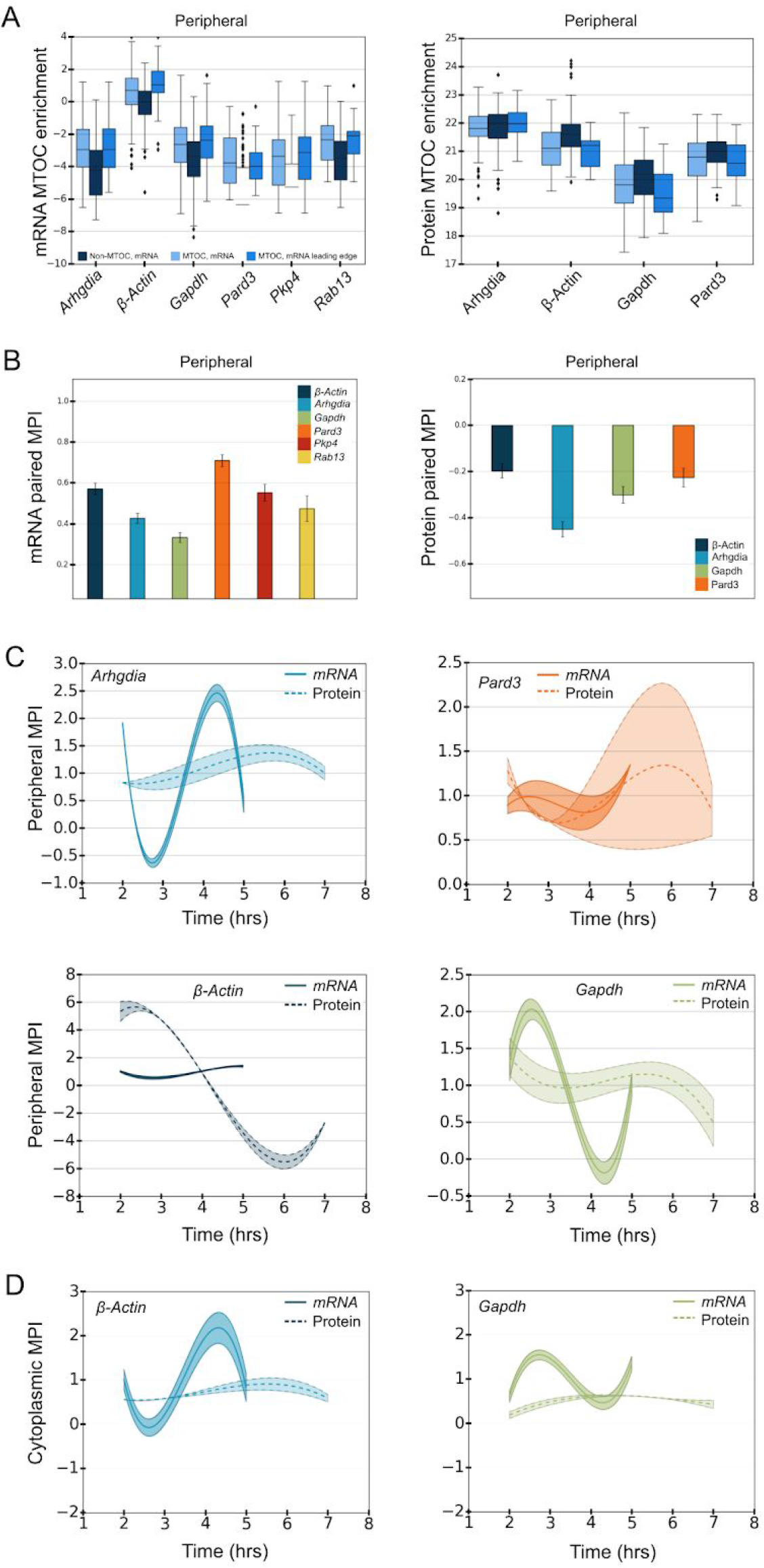
Correlative influence of peripheral mRNA and protein distributions and MTOC position. (A) The peripheral mRNA enrichment in non-MTOC containing quadrants, MTOC-containing quadrants and MTOC-containing quadrants when this quadrant coincides with the leading edge. The concentration of cytoplasmic *β-Actin*, *GAPDH* and *Rab13* transcripts and Arhgdia protein is enriched in the MTOC-containing quadrant when it is in the leading edge. (B) Comparison of MPI values for mRNAs and proteins in peripheral populations (all time points). (E) Comparison of MPI dynamics for β-Actin and GAPDH mRNA-protein pairs. Bar graphs in (B) show median and .25 and .75 quantile error bars for 11 bootstrapped MPI estimates.

**Figure S3, related to Figure 5.**
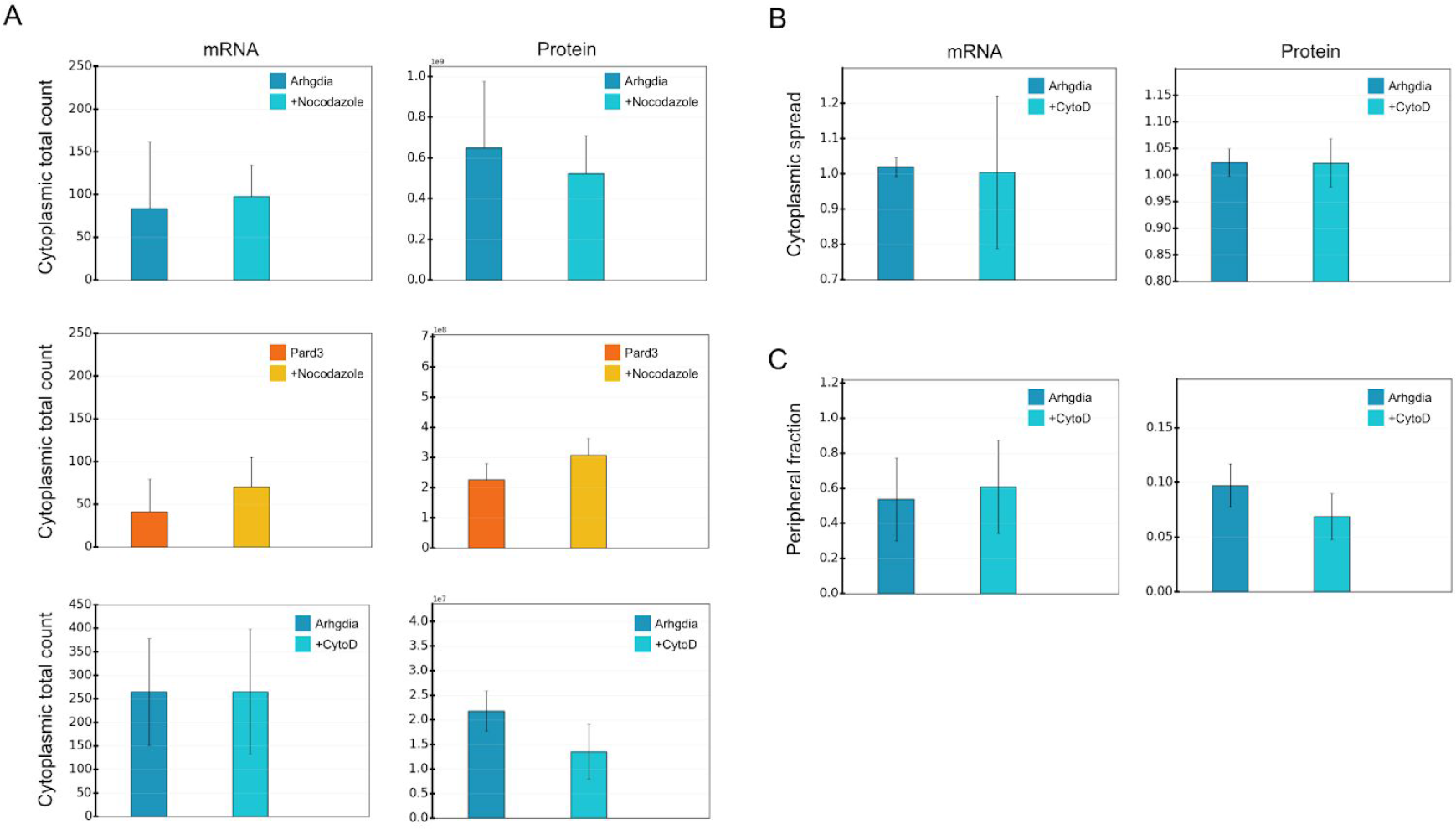
Effects of cytoskeleton disturbance on mRNA-protein cytoplasmic spread, counts and peripheral fraction. (A) + (B) Effects of Nocodazole and CytoD treatment on cytoplasmic total and cytoplasmic spread descriptors for Arhgdia and Pard3 mRNAs and proteins at 3-5 h time-points. Bar graphs show median with .25 and .75 quantile error bars. (C) The peripheral fraction of Arhgdia transcript and protein (at 3 and 5 h combined) were calculated similarly to Figure 5C.

**Figure S4, related to Figure 6.**
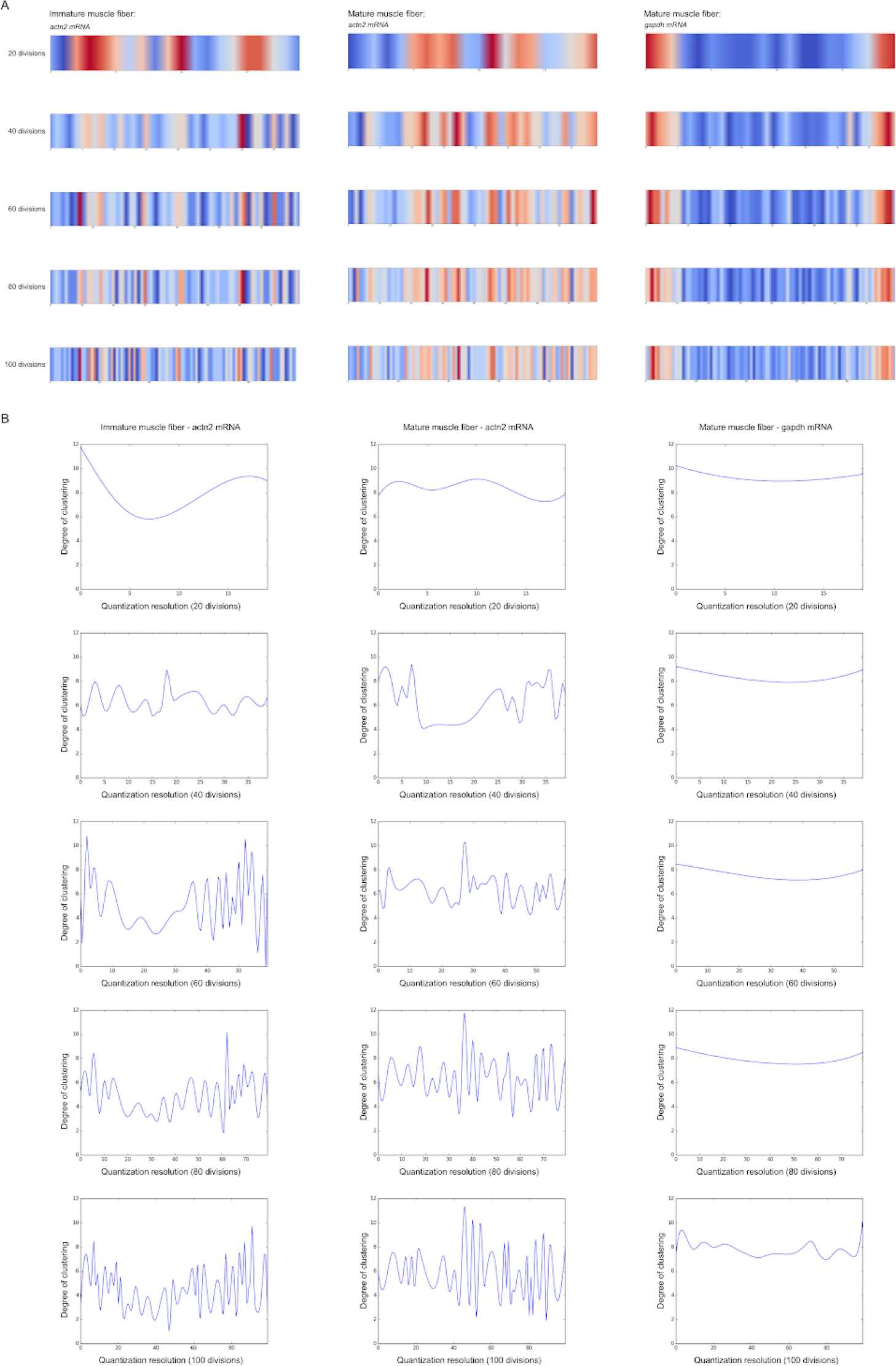
Sarcomeric mRNAs cluster in a striated pattern in differentiated myofibers. (A) + (B) The mRNA local density was computed between two nuclei. Each cell was quantized in vertical quadrants and relative concentration of mRNA in each quadrant was computed by normalizing the counts by the relevant surface. A wave-like clustering is observed for *actn2* in mature compared to immature fibers. No clustering is observed for *Gapdh*.

## Footnotes

1 Specific constants are dependent on the microscope and camera settings.

